# *Borrelia burgdorferi* PlzA is a cyclic-di-GMP dependent DNA and RNA binding protein

**DOI:** 10.1101/2023.01.30.526351

**Authors:** Nerina Jusufovic, Andrew C. Krusenstjerna, Christina R. Savage, Timothy C. Saylor, Catherine A. Brissette, Wolfram R. Zückert, Paula J. Schlax, Md A. Motaleb, Brian Stevenson

## Abstract

The PilZ domain-containing protein, PlzA, is the only known cyclic di-GMP binding protein encoded by all Lyme disease spirochetes. PlzA has been implicated in the regulation of many borrelial processes, but the effector mechanism of PlzA was not previously known. Here we report that PlzA can bind DNA and RNA and that nucleic acid binding requires c-di-GMP, with the affinity of PlzA for nucleic acids increasing as concentrations of c-di-GMP were increased. A mutant PlzA that is incapable of binding c-di-GMP did not bind to any tested nucleic acids. We also determined that PlzA interacts predominantly with the major groove of DNA and that sequence length plays a role in DNA binding affinity. PlzA is a dual-domain protein with a PilZ-like N-terminal domain linked to a canonical C-terminal PilZ domain. Dissection of the domains demonstrated that the separated N-terminal domain bound nucleic acids independently of c-di-GMP. The C-terminal domain, which includes the c-di-GMP binding motifs, did not bind nucleic acids under any tested conditions. Our data are supported by computational docking, which predicts that c-di-GMP binding at the C-terminal domain stabilizes the overall protein structure and facilitates PlzA-DNA interactions via residues in the N-terminal domain. Based on our data, we propose that levels of c-di-GMP during the various stages of the enzootic life cycle direct PlzA binding to regulatory targets.

## INTRODUCTION

Cyclic bis-(3’→5’)-dimeric guanosine monophosphate (c-di-GMP) is a ubiquitous bacterial cyclic di-nucleotide. In many bacterial species, it engages in the regulation of key processes such as transcription, motility, biofilm formation, and virulence (Ryjenkov *et al*., 2005; Hengge, 2009; Romling *et al*., 2013; Jenal *et al*., 2017; Valentini and Filloux, 2019). The Lyme disease spirochete, *Borrelia burgdorferi* sensu lato, referred to as *Borrelia burgdorferi* henceforth, has a c-di-GMP regulatory network consisting of a transmembrane sensor histidine kinase (Hk1), a response regulator diguanylate cyclase (Rrp1), two phosphodiesterases (PdeA and PdeB), and a single, chromosomally encoded PilZ domain-containing protein, PlzA (Galperin *et al*., 2001; Freedman *et al*., 2009; Rogers *et al*., 2009; Sultan *et al*., 2010; Caimano *et al*., 2011; Sultan *et al*., 2011; Novak *et al*., 2014).

The borrelial Hk1/Rrp1 two-component system synthesizes c-di-GMP; activation of Hk1 (BB_0420) occurs through an unknown signal (Caimano *et al*., 2011; Bauer *et al*., 2015). Rrp1 (BB_0419), the only identified diguanylate cyclase in *B. burgdorferi,* synthesizes c-di-GMP from two GTP molecules and is essential for acquisition and survival in the tick vector (Rogers *et al*., 2009; Kostick *et al*., 2011). Turnover of c-di-GMP is mediated by PdeA (BB_0363) and PdeB (BB_0374), the only phosphodiesterases present in *B. burgdorferi*, which degrade c-di-GMP to pGpG and GMP, respectively (Sultan *et al*., 2010; Sultan *et al*., 2011). *B. burgdorferi* must delicately balance c-di-GMP signaling, as c-di-GMP is required for both maintenance in the tick vector and successful transmission into the vertebrate host, but constitutive synthesis is detrimental to spirochete survival during vertebrate infection (Sultan *et al*., 2010; Sultan *et al*., 2011; Caimano *et al*., 2015; Groshong *et al*., 2021).

Synthesized c-di-GMP is bound by PlzA, the only universal c-di-GMP binding protein found in all isolates of Lyme disease *Borreliae* (Kostick-Dunn *et al*., 2018). Mutation of *plzA* has been implicated in many *B. burgdorferi* phenotypes, including dysregulation of alternative carbohydrate utilization, decreased motility, and virulence defects (Pitzer *et al*., 2011; He *et al*., 2014; Mallory *et al*., 2016; Zhang *et al*., 2018; Groshong *et al*., 2021). Binding of c-di-GMP induces conformational changes in PlzA structure, which has led to the hypothesis that c-di-GMP binding acts as a switching mechanism providing the holo and apo-forms of PlzA with distinct functions (Mallory *et al*., 2016; Groshong *et al*., 2021; Grassmann *et al*., 2023). Specifically, c-di-GMP bound PlzA is thought to regulate genes required for survival in the tick host, while apo-PlzA is speculated to play a role in regulation during vertebrate infection (Groshong *et al*., 2021; Grassmann *et al*., 2023).

A common mechanism of action for c-di-GMP effector proteins is to bind nucleic acids (Wilksch *et al*., 2011; Tschowri *et al*., 2014; Tan *et al*., 2015; Wang *et al*., 2016; Schäper *et al*., 2017; Hsieh *et al*., 2018; Skotnicka *et al*., 2020). It has been hypothesized that PlzA could bind DNA and regulate genes through impacts on transcription, such as directly interacting with RNA polymerase (RNAP), but this has yet to be validated experimentally (Groshong *et al*., 2021; Grassmann *et al*., 2023). Conversely, others have argued that PlzA cannot bind DNA, as it lacks an obvious, conventional DNA-binding motif (Caimano *et al*., 2015; Zhang *et al*., 2018). Indeed, Zhang *et al*. concluded that PlzA does not bind DNA, as their electrophoretic mobility shift assays (EMSA) did not identify binding to the *glp* promoter region (Zhang *et al*., 2018). We note that the raw data and methodologies, including protein and c-di-GMP concentrations, were not shown in that report. After the submission of a preprint of our manuscript, another group published that PlzA has RNA chaperone activity that is independent of c-di-GMP, supporting the notion that holo- and apo-forms of PlzA have distinct functions (Van Gundy *et al*., 2023).

Given the multiple conflicting hypotheses regarding PlzA function, as well as the potential for additional mechanisms between liganded and unliganded PlzA, we sought to independently characterize PlzA nucleic acid binding function. To that end, we chose a well-established locus PlzA is known to regulate, *glpFKD*, the glycerol catabolism operon. The *glpFKD* operon promotes *B. burgdorferi* survival in ticks (Pappas *et al*., 2011; Corona and Schwartz, 2015; Helble *et al*., 2021). After a tick has completed the digestion of its blood meal, it is hypothesized that the midgut is devoid of glucose and many other nutrients. To survive in that environment, Lyme disease spirochetes would need to utilize alternate carbon sources such as glycerol, which is available in the tick host (Lee and Baust, 1987). The *glpFKD* operon is regulated through the Hk1/Rrp1 pathway, and borrelial *plzA* mutants are defective in glycerol metabolism (Rogers *et al*., 2009; He *et al*., 2011; Caimano *et al*., 2015; Zhang *et al*., 2018). Further, PlzA was found to both positively and negatively affect the *glp* operon at higher and lower c-di-GMP levels, respectively (Zhang *et al*., 2018).

PlzA is classified as an xPilZ-PilZ domain protein, given its dual-domain structure. The binding of c-di-GMP to the interdomain linker and C-terminal PilZ domain results in a conformational change; locking the C-terminal PilZ and N-terminal xPilZ domains into rigid conformation, which we hypothesized could permit DNA binding (Mallory *et al*., 2016; Groshong *et al*., 2021; Singh *et al*., 2021). While no canonical DNA binding domain is evident in the PlzA amino acid sequence, our lab has discovered and characterized multiple nucleic acid binding proteins possessing novel binding motifs (Babb *et al*., 2006; Riley *et al*., 2009; Jutras *et al*., 2012; Jutras *et al*., 2013a; Jutras *et al*., 2013b; Zhang *et al*., 2018; Singh *et al*., 2021). Here, we report that PlzA can bind DNA and RNA directly downstream of the promoter and into the 5’ untranslated region (UTR) of the *glpFKD* operon and that binding to this region is c-di-GMP dependent.

## RESULTS

### PlzA binds *glpFKD* DNA and the interaction is c-di-GMP dependent

Previous studies demonstrated that c-di-GMP and PlzA modulate expression of the glycerol metabolism operon (Rogers *et al*., 2009; He *et al*., 2011; Caimano *et al*., 2015; Zhang *et al*., 2018). We hypothesized that PlzA might bind 5’ of the *glpFKD* operon. To that end, we sought to determine if PlzA could bind DNA between *glpFKD* and the upstream ORF BB_0239, which encodes a deoxyguanosine/deoxyadenosine kinase. The transcriptional boundaries and promoter elements of the *glpFKD* operon were previously mapped (Adams *et al*., 2017; Grove *et al*., 2017). A fluorescently labeled DNA fragment was generated by PCR amplification from a plasmid containing the 410 bp intergenic region between the upstream gene and the start of *glpF* (BB_0240) and is denoted as *glpFKD*(−219) (Figure 1).

**FIGURE 1.**
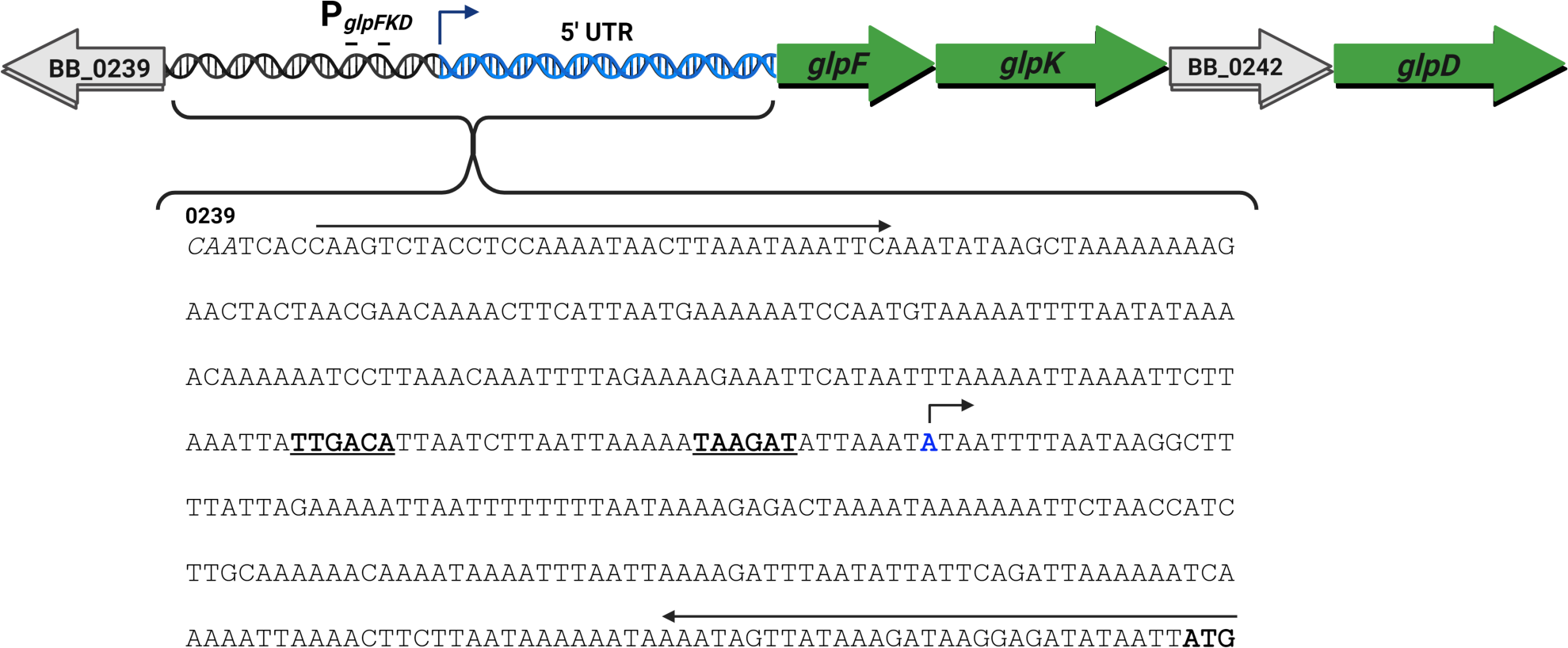
The *glpFKD* operon, 5’ UTR, and designed probe schematic. The glycerol catabolism gene operon (*glpFKD*) contains four genes: *glpF* (BB_0240, glycerol uptake facilitator protein), *glpK* (BB_0241, glycerol kinase), BB_0242 (hypothetical protein), and *glpD* (BB_0243, glycerol-3-phosphate dehydrogenase). The promoter and transcriptional boundaries were previously mapped (Grove *et al*., 2017). The −35 and −10 sites are bolded and underlined. The transcriptional start site nucleotide is highlighted in blue and topped with an arrow indicating the direction of transcription. The start of the *glpF* ORF is marked by the bolded ATG, while the start of the upstream BB_0239 ORF is labeled, and its nucleotides are italicized. The long arrows indicate the location of primers used to generate a large probe incorporating the entire intergenic region which is denoted as *glpFKD*(−219) for initial DNA binding studies.

To assess if PlzA could bind this DNA, EMSAs with increasing concentrations of recombinant PlzA were performed (Figure 2A). Binding was observed at higher micromolar concentrations, as evidenced by the disappearance of free DNA. The experiment was repeated with added non-specific competitor poly-dI-dC to a final concentration of 2.5 ng/µL. Poly-dI-dC did not appreciably compete away the shifted protein-DNA complexes (Figure 2B).

**FIGURE 2.**
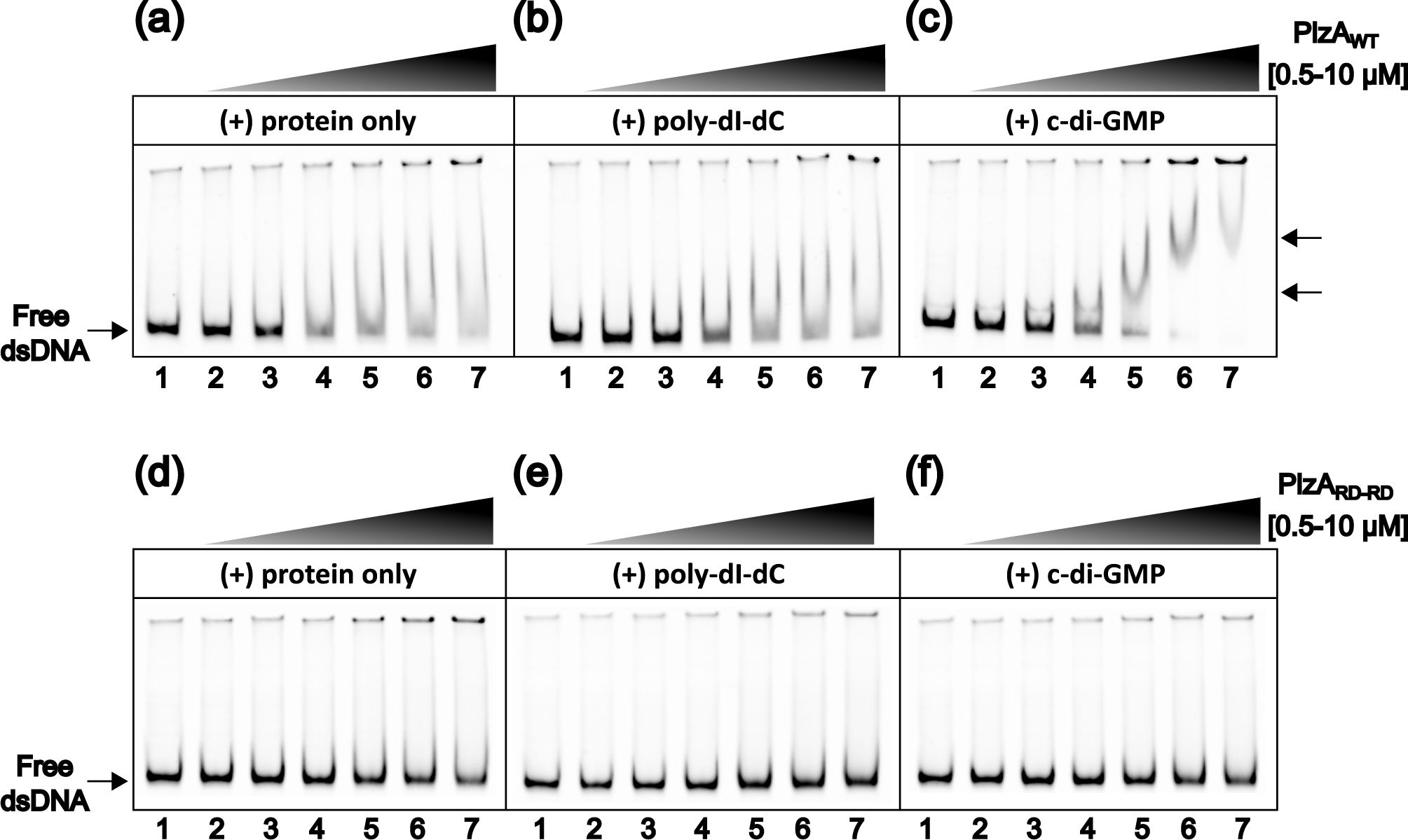
DNA binding by PlzA is c-di-GMP dependent. (**a**) EMSA with increasing concentrations of PlzA_WT_ showing binding to *glpFKD*(−219) DNA. The mixture was incubated for 10 min before electrophoresis. (**b**) The same EMSA mixture as in (**a**) but with a final concentration of 2.5 ng/µL of the nonspecific competitor poly-dI-dC. Protein was incubated with poly-dI-dC for 5 min prior to the addition of labeled *glpFKD*(−219) probe. (**c**) The same EMSA mixture as in (**b**) but supplemented with 100 µM c-di-GMP. Protein was incubated with c-di-GMP for 5 min prior to the addition of any nucleic acids. Arrows indicate PlzA-*glpFKD*(−219) complexes. Lane 1 in (**a-c**) is the probe only control, containing 50 nM of probe. Protein concentrations are as follows in each subsequent lane: Lane 2: 0.5 µM, Lane 3: 1 µM, Lane 4: 2.5 µM, Lane 5: 5 µM, Lane 6: 7.5 µM, and Lane 7: 10 µM. (**d**), (**e**), (**f**) EMSA experiments as in (**a**), (**b**), and (**c**), respectively, but performed with the mutant PlzA_RD-RD_ protein. All conditions were performed identically as with PlzA_WT_. Each panel is a representative gel from separate EMSAs which were performed in triplicate (PlzA_WT_) or duplicate (PlzA_RD-RD_).

Given that binding of c-di-GMP has been hypothesized to act as a switching mechanism for PlzA function, we next investigated the effects of c-di-GMP on DNA binding. EMSA reaction mixtures were supplemented with exogenous c-di-GMP to a final concentration of 100 µM. This concentration was used to saturate all PlzA molecules present in the reaction to ensure complete binding throughout electrophoresis. Saturating concentrations have similarly been used in other studies with c-di-GMP binding proteins that also bind DNA (Wilksch *et al*., 2011; Yang *et al*., 2013; Tan *et al*., 2015; Zamorano-Sánchez *et al*., 2015; Schäper *et al*., 2017; Skotnicka *et al*., 2020). The EMSA with *glpFKD*(−219), poly-dI-dC, and increasing PlzA was repeated with the supplemented c-di-GMP (Figure 2C). The addition of c-di-GMP enhanced PlzA binding affinity for *glpFKD*(−219), resulting in binding at lower concentrations of protein, and the formation of distinct protein-DNA complexes as indicated by the arrows. Further, as PlzA concentration increased, the mobility of the shifted band decreased.

To further assess the requirement of c-di-GMP for PlzA DNA binding activity, a mutant PlzA protein (PlzA_RD-RD_) that cannot bind c-di-GMP was produced by site-directed mutagenesis. This mutant replaced key arginine residues in the conserved c-di-GMP binding motif, RxxxR, with aspartic acids (Ryjenkov *et al*., 2006; Amikam and Galperin, 2006; Chou and Galperin, 2016). These mutations (R145D and R149D) have previously been shown to abolish c-di-GMP binding (Freedman *et al*., 2009; Mallory *et al*., 2016; Singh *et al*., 2021). The *glpFKD*(−219) EMSAs were repeated with PlzA_RD-RD_ (Figure 2D-F). Binding to *glpFKD*(−219) was not observed under any tested conditions with PlzA_RD-RD_.

To ensure binding was not due to the recognition of vector-derived sequences within the probe, competition EMSAs were performed with unlabeled DNA competitors derived from empty pCR 2.1 TA clones, or those containing the *glpFKD*(−219) sequence (Figure S1A). The nonspecific DNA from the empty pCR 2.1 vector did not compete away PlzA_WT_-*glpFKD*(−219) complexes at a 20-fold higher concentration (Figure S1B, lanes 3-5). Meanwhile, as little as 6-fold molar excess of unlabeled *glpFKD*(−219) was able to compete with the probe, and a 10-fold molar excess successfully competed away the PlzA_WT_-*glpFKD*(−219) complexes (Figure S1B, lanes 6-8). These results demonstrate PlzA preferentially bound the *glpFKD*-derived sequences.

Noting that c-di-GMP enhanced DNA binding by PlzA yet free DNA decreased with purified recombinant PlzA without adding exogenous c-di-GMP, we investigated the possibility that PlzA was being purified with c-di-GMP already bound. Purified recombinant proteins were prepared and assayed by liquid chromatography-tandem mass spectrometry. Extracted ion chromatograms showed the presence of c-di-GMP in PlzA_WT_ protein preparations but not in PlzA_RD-RD_ preparations (Figure 3). The concentration of c-di-GMP present in recombinant PlzA_WT_ was extrapolated from a c-di-GMP standard curve at 99.29 nM. A total of 31.7 µM of protein had been analyzed, indicating that approximately 0.3% of the protein was bound with c-di-GMP. Prior studies have reported dissociation constants of 1.25 to 6.25 µM of PlzA for c-di-GMP, which are typical K_D_ values for PilZ receptors and c-di-GMP (Pitzer *et al*., 2011; Mallory *et al*., 2016). The amount of co-purified c-di-GMP detected is lower than the K_D_. This is likely due to removal of some c-di-GMP during the protein purification process, as well as *E. coli* maintaining low cellular concentrations of c-di-GMP (∼40-80 nM) (Sarenko *et al*., 2017; Junkermeier and Hengge, 2023). Taken together, these data indicate that PlzA DNA binding to this region of the *glpFKD* operon is specific and c-di-GMP dependent.

**FIGURE 3.**
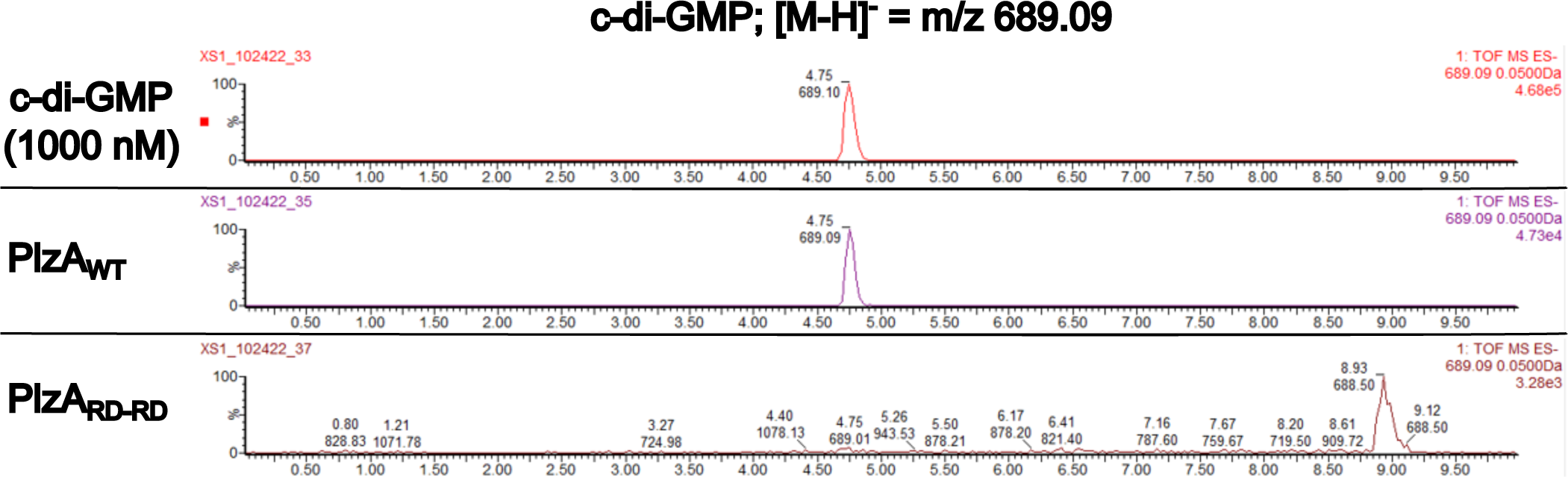
Detection of c-di-GMP in recombinant PlzA_WT_ and PlzA_RD-RD_ by LC-MS/MS. Recombinant PlzA_WT_ and PlzA_RD-RD_ were sent off for mass spectrometry to detect c-di-GMP. Extracted ion chromatograms of the LC-MS/MS analysis for detection of c-di-GMP are displayed. Chromatograms depict the control c-di-GMP standard at 1 µM, recombinant PlzA_WT_, and recombinant PlzA_RD-RD_. The LC-MS/MS analysis reveals PlzA_WT_ co-purified with c-di-GMP, but PlzA_RD-RD_ did not. The concentration of c-di-GMP in recombinant PlzA_WT_ was determined from a standard curve of c-di-GMP. PlzA_WT_ co-purified with 99.29 nM of c-di-GMP and approximately 0.3% of the protein was bound with c-di-GMP. Black lines indicate chromatograms unrelated to this figure that were removed.

### c-di-GMP levels modulate PlzA DNA binding affinity

It was previously identified with *glpFKD*-*gfp* promoter fusion studies that the minimal *glpFKD* promoter sequence (−46 relative to the *glpFKD* transcription start site) with the full 195bp UTR region was expressed at higher levels than the core promoter sequence alone (Grove *et al*., 2017). Given these findings and our data that PlzA can bind DNA, we focused further analysis of PlzA binding on a smaller sequence within that region that was found to be important for regulation (Grove *et al*., 2017; Savage *et al*., 2018). A 42 bp DNA probe encompassing the −7 to +35 sites, called *glpFKD*(−7/+35), was generated by annealing complementary oligonucleotides with the forward oligonucleotide 5’ conjugated to IRDye800 (Figure 4A). An RNA probe conjugated to Alexa Fluor 488 was also made for subsequent RNA binding studies.

**FIGURE 4.**
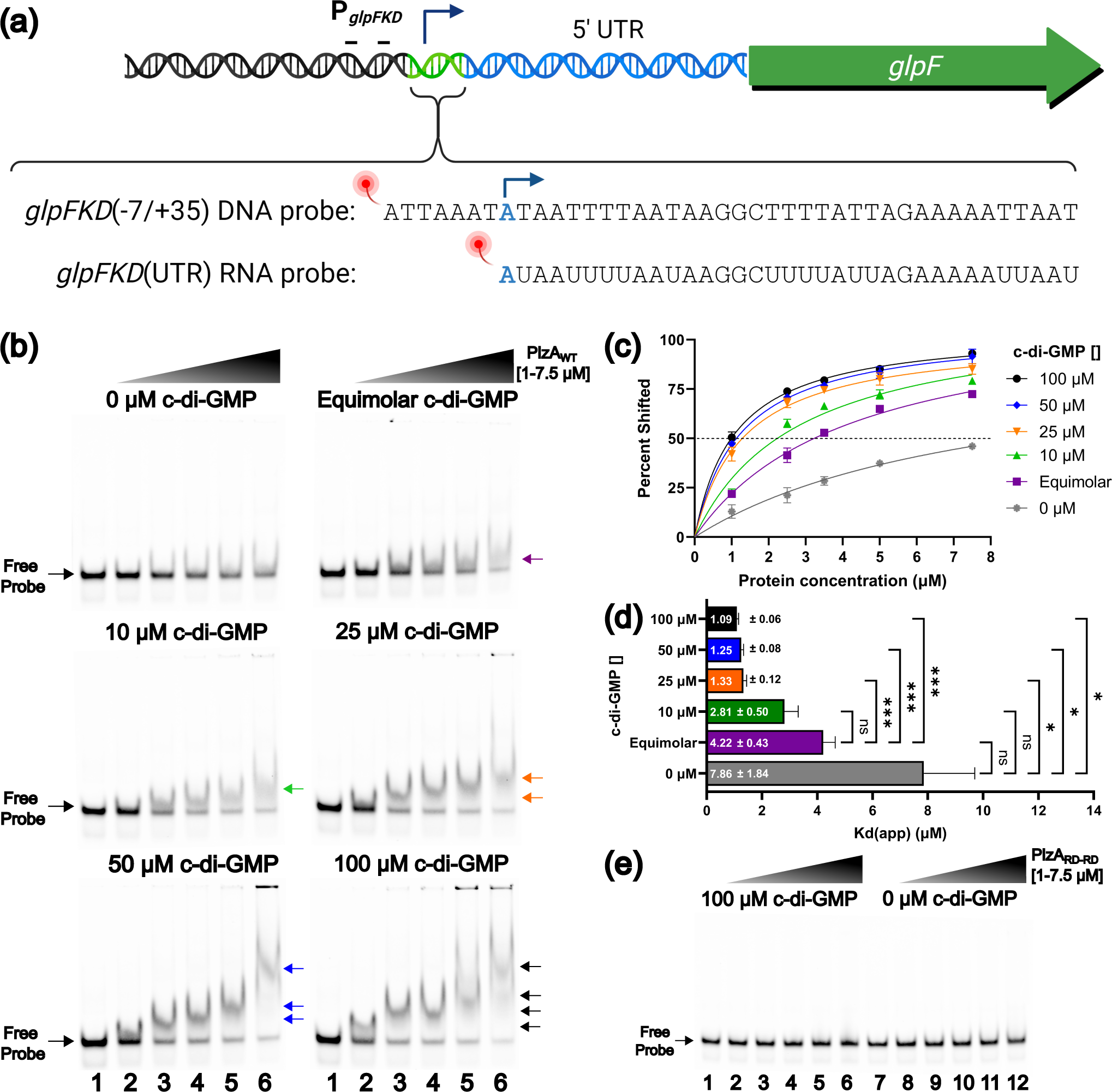
Levels of c-di-GMP alter DNA binding affinity of PlzA_WT_. (**a**) A diagram of the generated DNA probe, designated as *glpFKD*(−7/+35), corresponding to a 42 bp sequence adjacent to the −10 site and proceeding 35 bp into the 5’ UTR. An RNA probe was made corresponding to only nucleotides that are within the 5 ‘UTR which was designated as *glpFKD*(UTR) RNA. Probes were conjugated to a fluorescent molecule for the detection of binding in EMSAs. (**b**) EMSAs were performed with increasing protein concentrations of PlzA_WT,_ and 10 nM fluorescently labeled probe at varying c-di-GMP concentrations (0, equimolar, 10, 25, 50, or 100 µM). For each EMSA shown, the indicated c-di-GMP concentration was held constant in all lanes. For each EMSA shown, lanes 1 and 7 are probe-only controls. Protein concentrations were as follows: Lane 2-1 µM, Lane 3-2.5 µM, Lane 4-3.5 µM, Lane 5-5 µM, Lane 6-7.5 µM. A final concentration of 2.5 ng/µL of the nonspecific competitor poly-dI-dC was added to each lane. Arrows indicate higher-order complexes formed as protein concentration increases. A representative gel for each tested c-di-GMP concentration is shown from triplicate EMSAs. (**c**) Triplicate EMSAs were quantitated by densitometry to determine the K_d(app)_ of the PlzA_WT_-*glpFKD*(−7/35+) interaction by nonlinear regression analysis using the one-site specific binding setting in GraphPad Prism. Errors bars represent one standard deviation (SD). (**d**) Bar graph displaying the calculated K_d(app)_ ± the standard error of the mean (SEM). The K_d(app)_ values from three EMSAs were analyzed by a Welch ANOVA with Dunnett’s T3 multiple comparisons test via GraphPad Prism. Adjusted P-values are indicated as follows: ns= p >0.05, *= p ≤0.05, and ***= p ≤0.001. Comparisons between 10, 25, 50, and 100 µM c-di-GMP are not indicated on the graph as they were not significant. (**e**) A similar EMSA experiment was performed as in Figure 4B but with PlzA_RD-RD_ and either 100 (lanes 1-6) or 0 (lanes 7-12) µM of c-di-GMP supplemented. No binding is observed.

To determine how different c-di-GMP levels impact PlzA binding affinity, EMSAs with increasing PlzA_WT_ concentrations were performed at concentrations of added c-di-GMP ranging from 0 to 100 µM in the reaction mixtures with the *glpFKD*(−7/+35) probe (Figure 4B). Competitor poly-dI-dC was added at a final concentration of 2.5 ng/µL to reduce nonspecific interactions. PlzA_WT_ bound to *glpFKD*(−7/+35), and binding was enhanced by increasing the concentration of c-di-GMP. In reactions with no added c-di-GMP, we observed that the amount of free DNA decreased with increasing PlzA concentrations but did not detect a band corresponding to the protein-DNA complex. The decrease in free DNA with no added c-di-GMP is presumably due to the presence of co-purified c-di-GMP in recombinant PlzA. When equimolar c-di-GMP and protein levels were tested, a discernable shift (formation of a new band) was evident only at protein concentrations greater than 2.5 µM (purple arrow). As c-di-GMP concentrations were further increased, shifts became discernable at lower protein concentrations, with the strongest binding observed at c-di-GMP concentrations of 25, 50, and 100 µM. Further, as protein concentration was increased, we consistently observed progressive super-shifting of complexes in a dose-dependent manner (Figure 4B, colored arrows). At protein concentrations greater than or equal to 5 µM coupled with c-di-GMP concentrations greater than 25 µM, complexes appeared to smear, suggesting the observed super-shifting is due to complex dissociation. The stoichiometry of this complex(es) is under investigation.

Next, we calculated the apparent binding affinity (K_D(app)_) under these experimental conditions for PlzA for *glpFKD*(−7/+35) at the various c-di-GMP concentrations (Figure 4C). The calculations were performed under the assumption that a single molecule of PlzA binds to a single molecule of DNA. As noted above, it is possible that the protein-nucleic acid ratio is not 1:1, which could affect the calculated values. The values described below are the K_D(app)_ ± SEM (standard error of the mean). Without any supplemented c-di-GMP, the extrapolated K_D(app)_ of PlzA_WT_-*glpFKD*(−7/+35) was calculated as 7.86 (± 1.84 µM), however we again note that some c-di-GMP is already present in the PlzA_WT_ protein preparations (see above and Figure 3). Binding affinity increased approximately 1.5-fold under equimolar c-di-GMP and protein concentrations with a calculated K_D(app)_ of 4.22 (± 0.43 µM). Under saturating c-di-GMP conditions (exceeding the K_D_ of PlzA for c-di-GMP), binding affinity increased further. Supplementing binding reactions with 10 µM c-di-GMP improved PlzA-DNA binding 2.5-fold with a K_D(app)_ of 2.81 (± 0.50 µM). An approximately 6- to 7-fold increase in binding affinity was observed when the binding reaction was supplemented with either 25, 50, or 100 µM of c-di-GMP resulting in calculated K_D(app)s_ of 1.33 (± 0.12 µM), 1.25 (± 0.08 µM), and 1.09 µM (± 0.06 µM), respectively.

To determine if the calculated K_D(app)_ values differed significantly, Welch’s ANOVA followed by a Dunnett’s T3 multiple comparisons test was performed (Figure 4D). All calculated K_D(app)_ values differed significantly (p ≤0.05) from the 0 µM c-di-GMP K_D(app)_ except for the equimolar and 10 µM c-di-GMP conditions. Compared to the equimolar condition, the 25, 50, and 100 µM K_D(app)_ values differed significantly (p ≤0.001), while the 10 µM c-di-GMP K_D(app)_ value did not (p >0.05). When comparing the 10, 25, 50, and 100 µM K_D(app)_ values of PlzA_WT_-*glpFKD*(−7/+35) interaction, there were no significant differences (p >0.05). The mutant PlzA_RD-RD_ did not bind *glpFKD*(−7/+35) under any tested conditions (Figure 4E). Overall, we conclude that levels of c-di-GMP significantly impact PlzA-DNA binding.

### PlzA interacts predominantly with the major groove of DNA and sequence length appears to play a role in DNA binding affinity

We next sought to gain further information on the mode of binding between PlzA and dsDNA. To explore these interactions, we performed binding assays with methyl green and actinomycin D which bind the major and minor grooves of DNA, respectively (Kim and Nordén, 1993; Basu *et al*., 2009; Liu *et al*., 2016; Shi *et al*., 2022). EMSAs were performed with increasing concentrations of these drugs to determine if either could compete with PlzA_WT_ for binding to the *glpFKD*(−7/+35) probe (Figure 5). Methyl green was able to compete for binding as evidenced by the increase in free DNA and by the disruption of PlzA-*glpFKD*(−7/+35) complex formation at higher concentrations. Oppositely, competition with actinomycin D was only evident at the highest concentrations tested. These results indicate that PlzA interacts with dsDNA mainly through interactions with the major groove.

**FIGURE 5.**
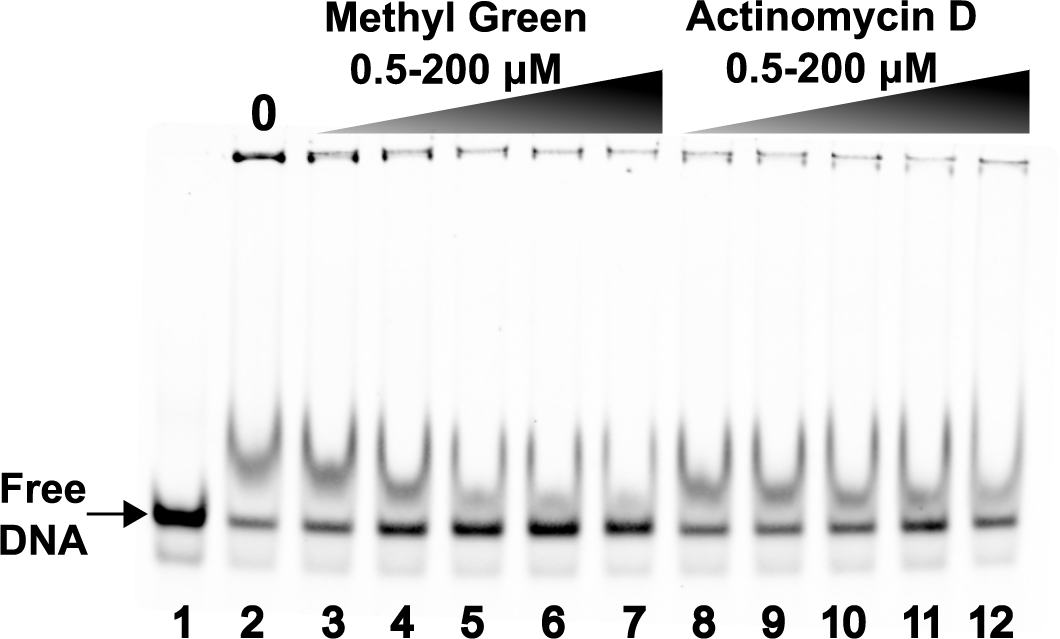
Major and Minor Groove Binding Assay. Competition EMSAs were performed with a constant PlzA_WT_ concentration of 3.5 µM, 10 nM fluorescently labeled *glpFKD*(−7/35+) probe, and a constant c-di-GMP concentration of 100 µM. Lane 1 is the probe-only control, while lane 2 is the probe + protein only control. Lanes 3-7 contain increasing concentrations of methyl green: Lane 3-0.5 µM, Lane 4-5 µM, Lane 5-50 µM, Lane 6-3 100 µM, and Lane 7-200 µM. Lanes 8-12 contain increasing concentrations of Actinomycin D: Lane 8-0.5 µM, Lane 9-5 µM, Lane 10-50 µM, Lane 11-100 µM and Lane 12-200 µM. A representative gel is shown from three independently performed experiments.

To gain insights on the DNA sequence requirements for PlzA binding, the *glpFKD*(−7/+35) sequence was changed at several nucleotides (Figure 6A). Mutant competitors 1 and 2 targeted the middle region of the sequence containing one of the few GC rich regions. Mutant competitors 3-5 swapped an ATTA repeat to CGGC at the 5’, 3’ or both ends to introduce greater GC content and potentially disrupt symmetry. Mutant competitors 7-9 were designed to disrupt potential secondary structural elements. Competition EMSAs with the IRDye labeled *glpFKD*(−7/+35) probe and the unlabeled mutagenized competitors were performed with PlzA_WT_ (Figure 6B-E). Despite introducing various mutations into the *glpFKD*(−7/+35) sequence, competition still occurred, although some competed slightly better than others. For example, mutant sequence 4, altering an ATTA to CGGC at the 5’ end, disrupted complex formation less than altering an ATTA at the 3’ end or mutating both. Our results suggest that the *glpFKD*(−7/+35) sequence may be a non-consensus or low affinity DNA binding site of PlzA within the *glpFKD* operon or that PlzA binds in a sequence independent manner. This 42 bp sequence was chosen as it is crucial for *glpFKD* regulation and is known to be bound by other proteins (Grove *et al*., 2017; Savage *et al*., 2018; Saylor *et al*., 2023; Zhang *et al*., 2024). We hypothesize that there could be higher affinity binding sites upstream of this operon and within the *B. burgdorferi* genome.

**FIGURE 6.**
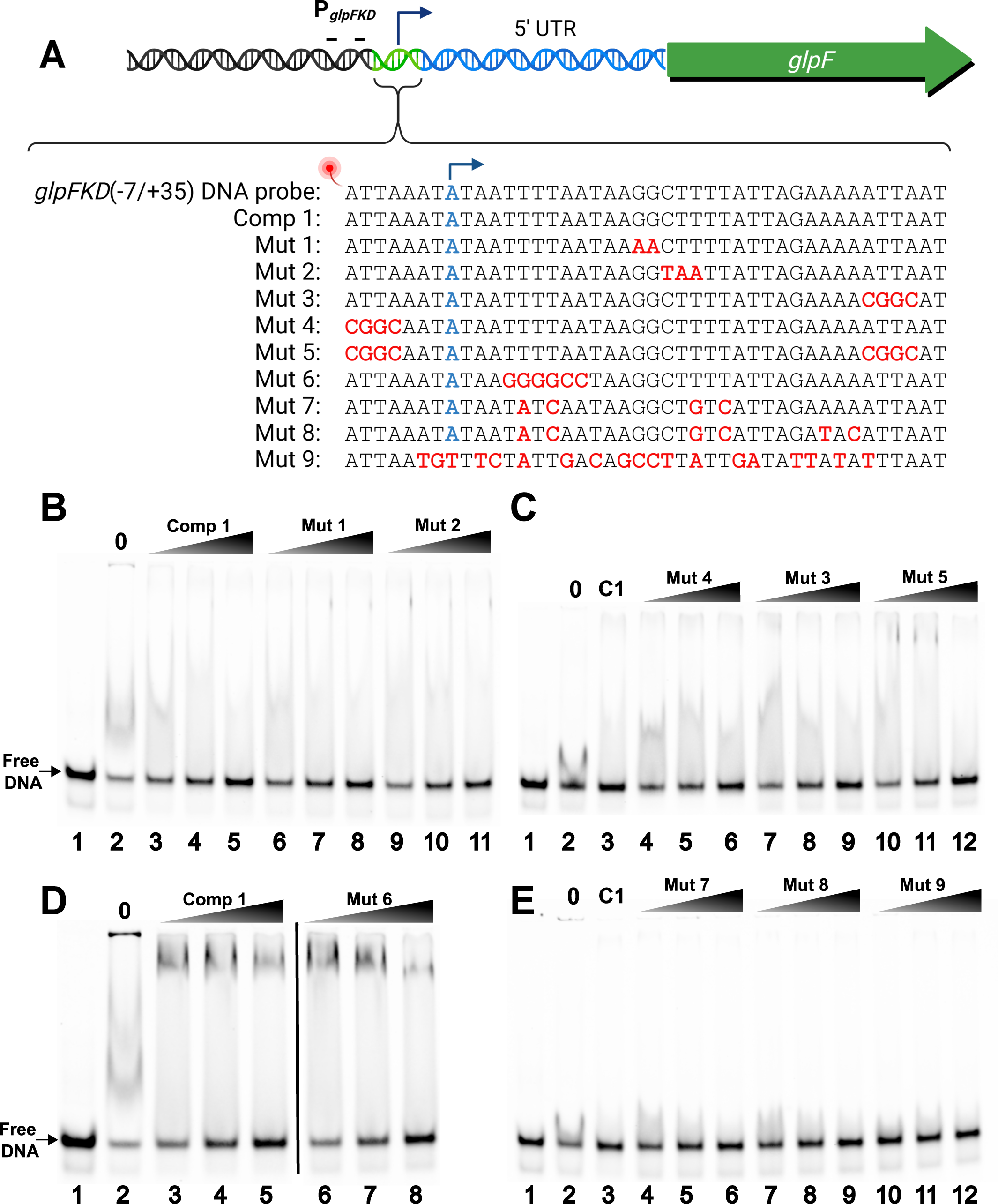
Competition EMSAs with mutagenized *glpFKD*(−7/35+) competitors. (**a**) A diagram of the generated unlabeled, mutagenized *glpFKD*(−7/+35) competitors. Comp 1 (C1) is the unlabeled, specific *glpFKD*(−7/+35) competitor. Introduced mutations are indicated in red for each mutant competitor. The transcriptional start site is the nucleotide labeled as blue. Competition EMSAs were performed with the various mutant competitors and the *glpFKD*(−7/35+) probe. (**b**) PlzA_WT_, c-di-GMP, poly-dI-dC, and *glpFKD*(−7/35+) probe concentrations were held constant at 2.5 µM, 100 µM, 2.5 ng/µL, and 10 nM respectively. Lane 1 is the probe-only control, while lane 2 is the probe + protein only control. Lanes 3-5: 250, 500, & 1000x molar excess (relative to the probe) of comp 1 (unlabeled, *glpFKD*(−7/35+). Lanes 6-8: Lanes 3-5: 250, 500, & 1000x mut 1 competitor. Lanes 6-8: Lanes 9-11: 250, 500, & 1000x mut 2 competitor. (**c**) Concentrations for protein, effector, nonspecific competitor, and probe as in (**a**). Lane 1 is the probe-only control, while lane 2 is the probe + protein only control. Lane 3 is the specific competitor control with 1000x of comp 1. Lanes 4-6: 250, 500, & 1000x mut 4 competitor. Lanes 7-9: 250, 500, & 1000x mut 3 competitor. Lanes 10-12: 250, 500, & 1000x mut 5 competitor. (**d**) PlzA_WT_, c-di-GMP, poly-dI-dC, and *glpFKD*(−7/35+) probe concentrations were held constant at 3.5 µM, 100 µM, 2.5 ng/µL, and 10 nM respectively. Lane 1 is the probe-only control, while lane 2 is the probe + protein only control. Lanes 3-5: 250, 500, & 1000x molar excess (relative to the probe) of comp 1 (unlabeled, *glpFKD*(−7/35+). Lanes 6-8: Lanes 3-5: 250, 500, & 1000x mut 6 competitor. The black line denotes irrelevant lanes that were removed from the same gel. (**e**) PlzA_WT_, c-di-GMP, poly-dI-dC, and *glpFKD*(−7/35+) probe concentrations were held constant at 2.0 µM, 100 µM, 2.5 ng/µL, and 10 nM respectively. Lane 1 is the probe-only control, while lane 2 is the probe + protein only control. Lane 3 is the specific competitor control with 1000x of comp 1. Lanes 4-6: 250, 500, & 1000x mut 7 competitor. Lanes 7-9: 250, 500, & 1000x mut 8 competitor. Lanes 10-12: 250, 500, & 1000x mut 9 competitor.

Since we could not identify an obvious PlzA sequence motif, we next investigated if PlzA bound DNA in a length-dependent mechanism. The *glpFKD*(−7/+35) sequence was truncated by nine bases at the 5’ end and then an additional three nucleotides at a time, and competition EMSAs were performed with these DNAs (Figure 7A). The truncated *glpFKD*(−7/+35) competitors exhibited less competition than the unlabeled specific competitor (Comp 1/C1) at the same concentration (Figure 7B). This could indicate that the nucleotides removed from the 5’ end are necessary for PlzA recognition or that PlzA has a higher affinity for longer pieces of DNA. Studies are ongoing to determine how PlzA recognizes nucleic acids and to find additional PlzA binding sites, as further elaborated upon in the discussion.

**FIGURE 7.**
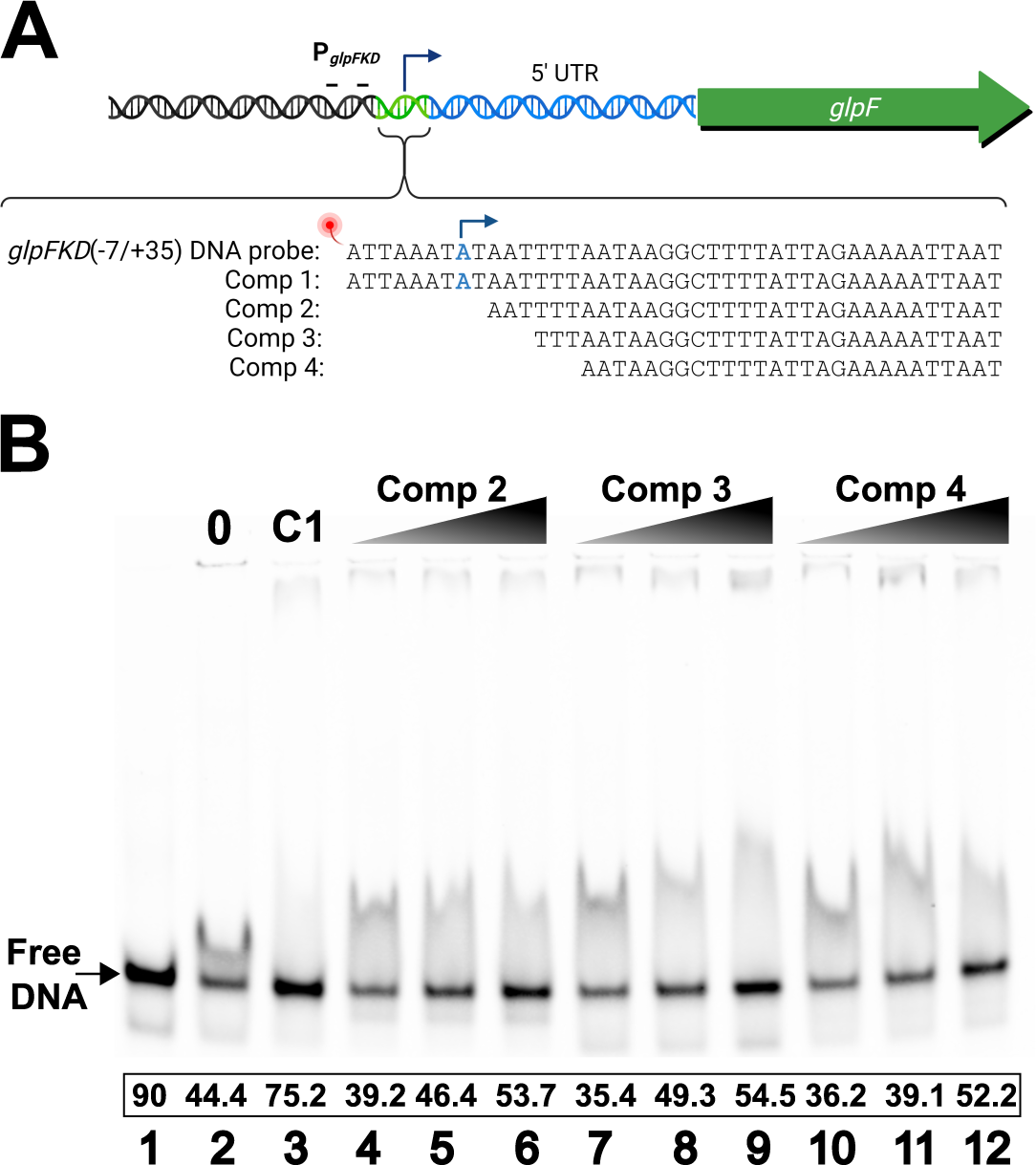
Competition EMSAs with truncated *glpFKD*(−7/35+) competitors. (**a**) A diagram of the generated unlabeled, truncated *glpFKD*(−7/+35) competitors. Comp 1 (C1) is the unlabeled, specific *glpFKD*(−7/+35) competitor. Competitors 2, 3, and 4 are unlabeled *glpFKD*(−7/+35) truncated at the 5’ end by either 9, 12, or 15 bases respectively. The transcriptional start site is the nucleotide labeled as blue. (**b**) PlzA_WT_, c-di-GMP, poly-dI-dC, and *glpFKD*(−7/35+) probe concentrations were held constant at 2.0 µM, 100 µM, 2.5 ng/µL, and 10 nM respectively. Lane 1 is the probe-only control, while lane 2 is the probe + protein only control. Lane 3 is the specific competitor control with 1000x of comp 1. Lanes 4-6: 250, 500, & 1000x comp 2 competitor. Lanes 7-9: 250, 500, & 1000x comp 3 competitor. Lanes 10-12: 250, 500, & 1000x comp 4 competitor. A representative gel is shown from two independently performed experiments.

### PlzA also binds *glp*-derived single-stranded nucleic acids

Several borrelial proteins that were initially identified as having affinities for DNA have subsequently been found to also bind RNA (Jutras *et al*., 2013a; Savage *et al*., 2018; Krusenstjerna *et al*., 2023). Given the role the 5’ UTR of the *glpFKD* operon plays in its regulation, we hypothesized PlzA could potentially also bind RNA. RNA probes used were designed to correspond to only transcribed regions (Figure 4A). The *glpFKD*(−7/+35) counterpart RNA probe is designated *glpFKD*(UTR). EMSAs were performed with increasing concentrations of PlzA_WT_ with 0 µM, 100 µM, or equimolar c-di-GMP levels and labeled *glpFKD*(UTR) (Figure 8). PlzA_WT_ bound the RNA, which was enhanced by addition of c-di-GMP (Figure 8A). The binding pattern to the *glpFKD*(UTR) RNA was comparable to the *glpFKD*(−7/+35) DNA, with increased protein and c-di-GMP concentrations leading to an increased size of the PlzA-RNA shifts.

**FIGURE 8.**
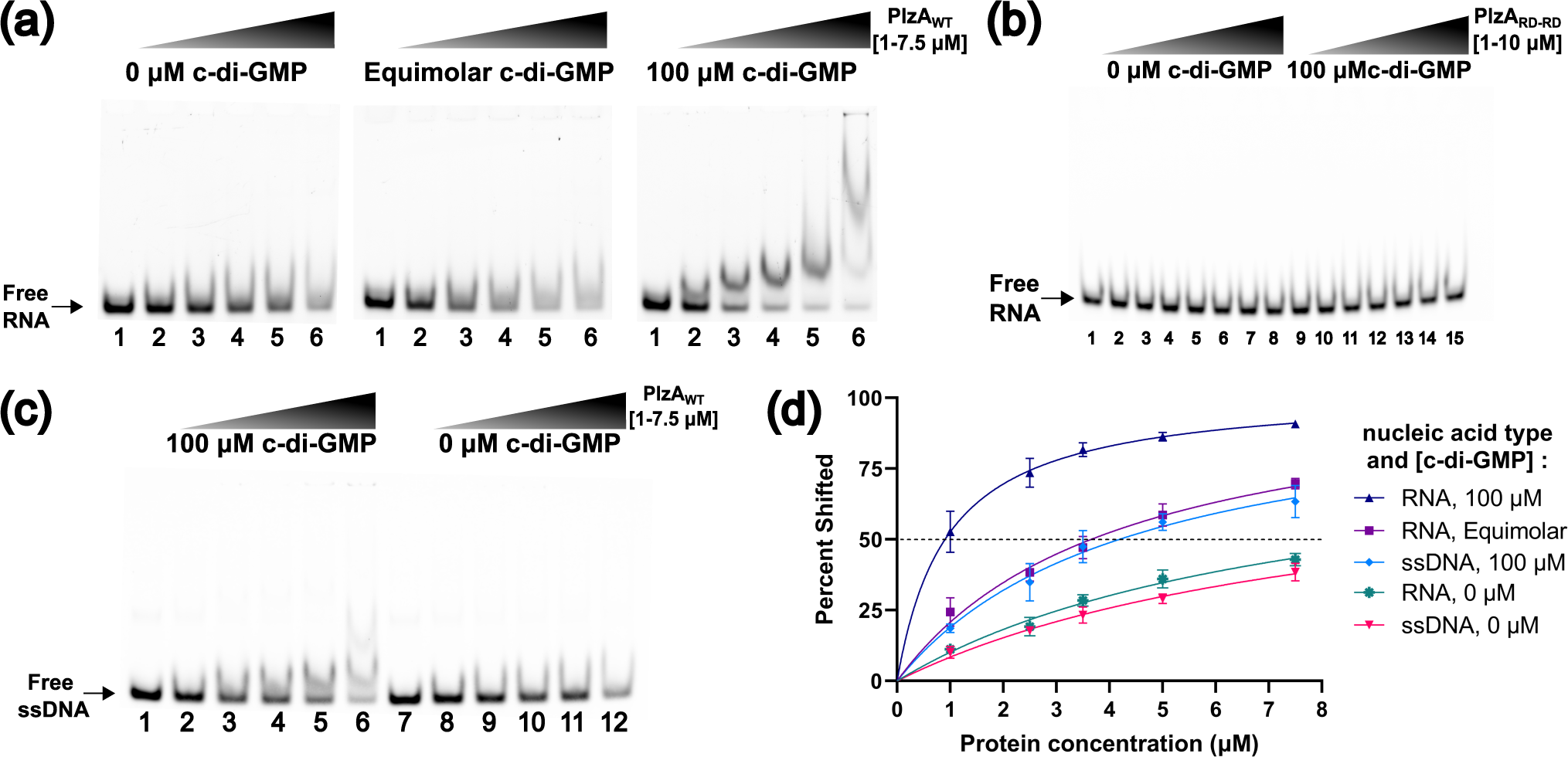
PlzA_WT_ also binds single stranded RNA and DNA. (**a**) Gradient EMSAs were performed with increasing protein concentrations of PlzA_WT_ without c-di-GMP (left panel), equimolar (middle panel), or 100 µM c-di-GMP (right panel) and 10 nM fluorescently labeled *glpFKD*(UTR) RNA probe. Lane 1 is the probe-only control in each EMSA. Lane protein concentrations were as follows: Lane 2-1 µM, Lane 3-2.5 µM, Lane 4-3.5 µM, Lane 5-5 µM, Lane 6-7.5 µM. (**b**) A similar EMSA experiment was performed with PlzA_RD-RD_ with (lanes 1-8) or without (lanes 9-15) 100 µM c-di-GMP and 50 nM of *glpFKD*(UTR) RNA probe. Lane 1 is a probe-only control. Protein concentrations were as follows: Lanes 2 and 9-1 µM, Lanes 3 and 10-1.25 µM, Lanes 4 and 11-2 µM, Lanes 5 and 12-2.5 µM, Lanes 6 and 13-3.3 µM, Lanes 7 and 14-5 µM, Lanes 8 and 15-10 µM. Riboguard was supplemented in the PlzA_RD-RD_ EMSA mixtures to a final concentration of 4 U/µL. (**c**) Gradient EMSAs were performed with increasing protein concentrations of PlzA_WT_ with 100 µM c-di-GMP (left panel) or 0 µM c-di-GMP (right panel) with 10 nM fluorescently labeled ss*glpF* probe. Lanes 1 and 7 are the probe-only control in each EMSA. Lane protein concentrations were as follows: Lanes 2 and 8-1 µM, Lanes 3 and 9-2.5 µM, Lanes 4 and 10-3.5 µM, Lanes 5 and 11-5 µM, Lanes 6 and 12-7.5 µM. Representative gels are shown from triplicate (PlzA_WT_) or duplicate (PlzA_RD-RD_) EMSA experiments. (**d**) Triplicate EMSAs from (**a**) and (**c**) were quantitated by densitometry to determine the K_d(app)_ of PlzA_WT_ for the *glpFKD* RNA or ssDNA interactions by nonlinear regression analysis using the one-site specific binding setting in GraphPad Prism. Errors bars represent one standard deviation (SD).

EMSAs were repeated with the PlzA_RD-RD_ mutant protein at 0 or 100 µM c-di-GMP (Figure 8B). The PlzA_RD-RD_ protein did not bind *glpFKD*(UTR) RNA at any tested protein concentration, regardless of whether c-di-GMP was included.

Given that PlzA can bind to both the DNA and RNA, we sought to determine if it could also bind the cognate ssDNA corresponding to the coding strand. This is relevant as ssDNA is exposed at open transcriptional complexes. The forward, IRDye labeled oligonucleotide used to generate the *glpFKD*(−7/+35) dsDNA probe was used as a single-stranded DNA probe (ss*glpF*) in EMSAs. No evident complexes were formed when exogenous c-di-GMP was excluded from EMSA reactions, but free ssDNA did decrease at the highest tested PlzA concentrations of 3.5-7.5 µM (Figure 8C right panel). Adding 100 µM c-di-GMP resulted in the formation of PlzA-ssDNA complexes but at higher concentrations of protein than required to achieve complex formation with dsDNA and RNA (Figure 8C left panel).

We next quantified the binding affinity of PlzA for the *glp* derived RNA and ssDNA sequences through densitometry of the EMSA shifts, and the apparent K_D(app)s_ calculated as described above (Figure 4D). As with dsDNA, the binding affinity of PlzA for RNA was weakest when no exogenous c-di-GMP was supplemented with a calculated K_D(app)_ of 7.64 (± 1.85 µM). Binding improved approximately 2-fold in equimolar c-di-GMP conditions with a K_D(app)_ of 4.14 (± 0.74 µM). The K_D(app)_ was 0.95 (± 0.11 µM) when the c-di-GMP concentration was 100 µM, an 8-fold increase in PlzA binding affinity for the *glpFKD*(UTR) RNA as compared to when no c-di-GMP was added. There was a significant difference between the PlzA_WT_-*glpFKD*(UTR) RNA K_D(app)_ value calculated at 100 µM c-di-GMP and both the equimolar and 0 µM c-di-GMP values (p ≤0.01) (Figure 9A). No significant differences were detected between the K_D(app)_ values calculated at 0 µM and equimolar c-di-GMP levels (p >0.05).

**FIGURE 9.**
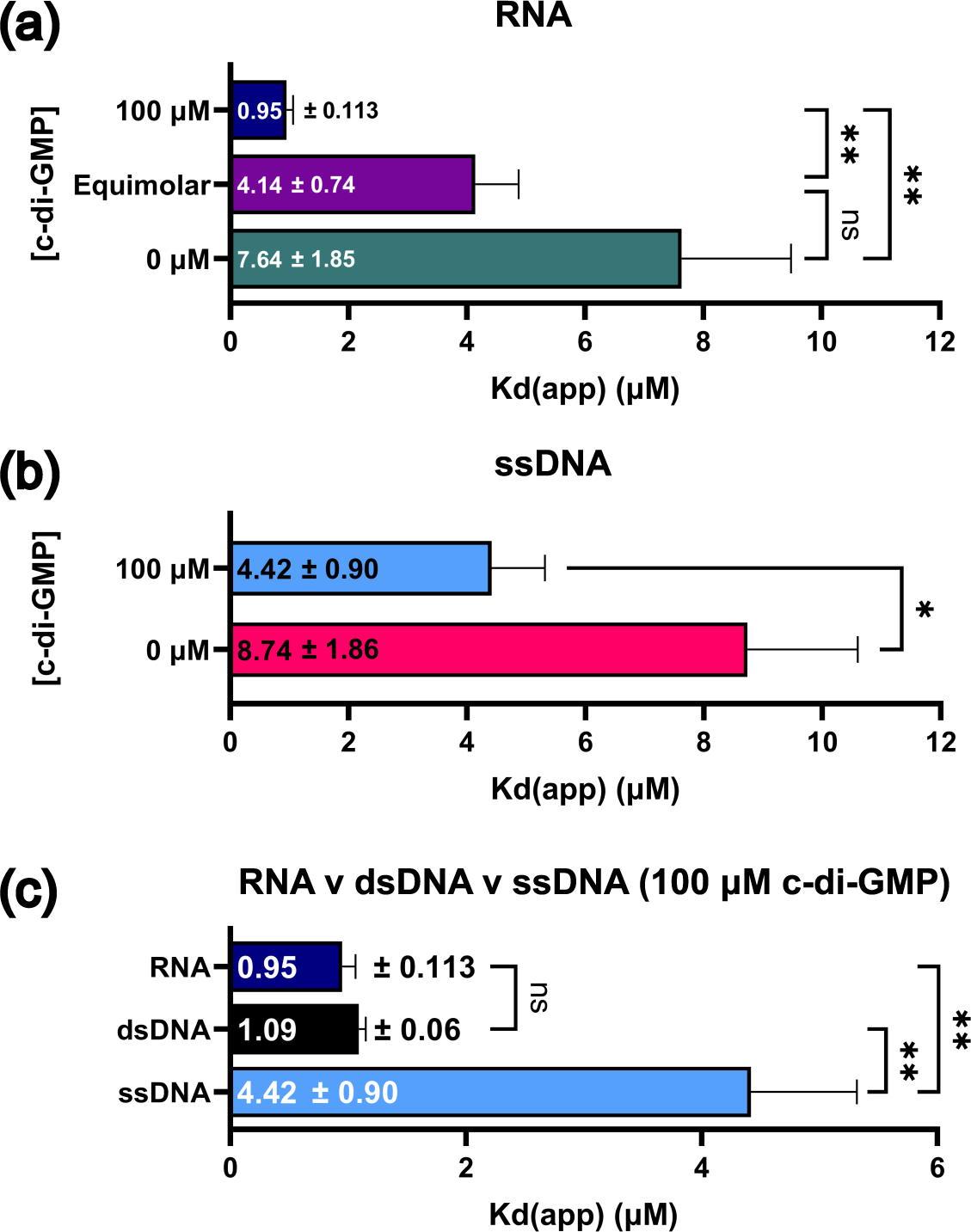
c-di-GMP significantly impacts PlzA-nucleic acid binding and PlzA prefers dsDNA and RNA over ssDNA. (**a**) The K_d(app)_ values from triplicate RNA EMSAs at each tested c-di-GMP concentration in Figure 5A were analyzed by a Welch ANOVA with Dunnett’s T3 multiple comparisons test. (**b**) The K_d(app)_ values from triplicate ssDNA EMSAs at the tested c-di-GMP concentrations in Figure 5C were analyzed by a Welch’s t-test. (**c**) The K_d(app)_ values from EMSAs with PlzA and each tested nucleic acid at 100 µM c-di-GMP were compared via a Welch ANOVA with Dunnett’s T3 multiple comparisons test. Bar graphs display the calculated K_d(app)s_ ± the standard error of the mean (SEM). Adjusted P-values are indicated as follows: ns= p >0.05, *= p ≤0.05, and **= p ≤0.01.

Although PlzA bound the ss*glpF* probe, it had a substantially lower affinity for ssDNA than dsDNA or RNA (Figure 8D). The K_D(app)_ of the PlzA_WT_-ss*glpF* interaction was 8.74 (± 1.86 µM) with 0 µM c-di-GMP and 4.42 (± 0.90 µM) with 100 µM c-di-GMP. These values were found to be statistically significant from one another (p <0.05) (Figure 9B). Only a 2-fold increase in binding affinity of PlzA for the single-strand DNA was observed when the c-di-GMP concentration was increased to 100 µM, whereas the same concentration of c-di-GMP increased PlzA binding affinity 7-fold for dsDNA and 8-fold for RNA.

Lastly, we also compared the binding affinities of PlzA for all three tested nucleic acids (Figure 9C). The calculated K_D(app)_ values of PlzA_WT_ for each nucleic acid in the presence of 100 µM c-di-GMP were compared with a Welch’s ANOVA and Dunnett’s T3 multiple comparisons test. No significant difference (p >0.05) was detected between the PlzA K_D(app)_ values for dsDNA (1.09 ± 0.06 µM) and RNA (0.95 ± 0.11 µM), indicating that PlzA binds the *glpFKD* dsDNA and RNA with similar affinity. Both the dsDNA and RNA K_D(app)s_ were significantly different from the calculated ssDNA K_D(app)_ (4.42 ± 0.90 µM) (p ≤0.01), suggesting that PlzA prefers *glp*-derived dsDNA and RNA over ssDNA. All the binding data presented throughout the manuscript are summarized in Table 1. Collectively, our data indicate that PlzA binding activity is c-di-GMP dependent for all tested *glpFKD* nucleic acids.

**Table 1.** Binding interaction summary of PlzA_WT_ and the tested nucleic acids at various c-di-GMP concentrations.

| *glpFKD(-7/+35)* dsDNA |  |  |  |
| --- | --- | --- | --- |
| c-di-GMP (μM) | K<sub>d(app)</sub> (μM), 95% C.I. [LL, UL] | B<sub>max</sub> (% Shifted), 95% C.I. [LL, UL] | Adj. R-squared |
| 100 | 1.09, [0.97, 1.23] | 105.20, [102.20, 108.40] | 0.986 |
| 50 | 1.25, [1.08, 1.45] | 105.50, [101.50, 109.90] | 0.981 |
| 25 | 1.33, [1.10, 1.59] | 101.60, [96.58, 107.10] | 0.971 |
| 10 | 2.81, [1.98, 4.02] | 113.00, [98.79, 132.30] | 0.940 |
| Equimolar | 4.22, [3.40, 5.26] | 115.40, [104.40, 129.10] | 0.983 |
| 0 | 7.86, [4.85, 14.06] | 94.25, [71.72, 138.80] | 0.950 |
| *glpFKD(UTR)* RNA |  |  |  |
| c-di-GMP (μM) | K<sub>d(app)</sub> (μM), 95% C.I. [LL, UL] | B<sub>max</sub> (% Shifted), 95% C.I. [LL, UL] | Adj. R-squared |
| 100 | 0.95, [0.73, 1.22] | 102.6, [96.76, 109.20] | 0.930 |
| Equimolar | 4.14, [2.79, 6.30] | 106.7, [89.50, 132.60] | 0.944 |
| 0 | 7.64, [4.71, 13.66] | 87.73, [66.88, 128.80] | 0.950 |
| *glpF* ssDNA |  |  |  |
| c-di-GMP (μM) | K<sub>d(app)</sub> (μM), 95% C.I. [LL, UL] | B<sub>max</sub> (% Shifted), 95% C.I. [LL, UL] | Adj. R-squared |
| 100 | 4.42, [2.94, 6.84] | 102.80, [85.32, 129.90] | 0.920 |
| 0 | 8.74, [5.67, 14.70] | 82.04, [63.53, 116.90] | 0.950 |

### Dissection of PlzA reveals the nucleic acid binding domain

The first discovered c-di-GMP binding receptors were the PilZ domain-containing proteins (Ryjenkov *et al*., 2006; Amikam and Galperin, 2006; Galperin and Chou, 2020). The PilZ domain consists of two conserved c-di-GMP binding motifs, RxxxR and [D/N]ZSXXG (Ryjenkov *et al*., 2006; Amikam and Galperin, 2006; Chou and Galperin, 2016; Cheang *et al*., 2019). The recently solved crystal structure of *B. burgdorferi* PlzA revealed a unique dual-domain topology consisting of an amino-terminal PilZ-like domain, called PilZN3, which is connected to the carboxy-terminal PilZ domain via a linker domain that contains the RxxxR c-di-GMP binding motif (Singh *et al*., 2021). The first 141 amino acid residues make up the N-terminus of PlzA (PlzA_NTD_), while residues 142-261 make up the C-terminal (PlzA_CTD_) PilZ domain. Having determined that PlzA is a c-di-GMP-dependent DNA and RNA binding protein, we next sought to determine which domain of PlzA binds nucleic acid. To generate recombinant PlzA_NTD_ and PlzA_CTD_ domains, truncated *plzA* genes called *plzA*-NTD and *plzA*-CTD were cloned to encode the residues described above, which correspond to each respective domain (Figure 10A).

**FIGURE 10.**
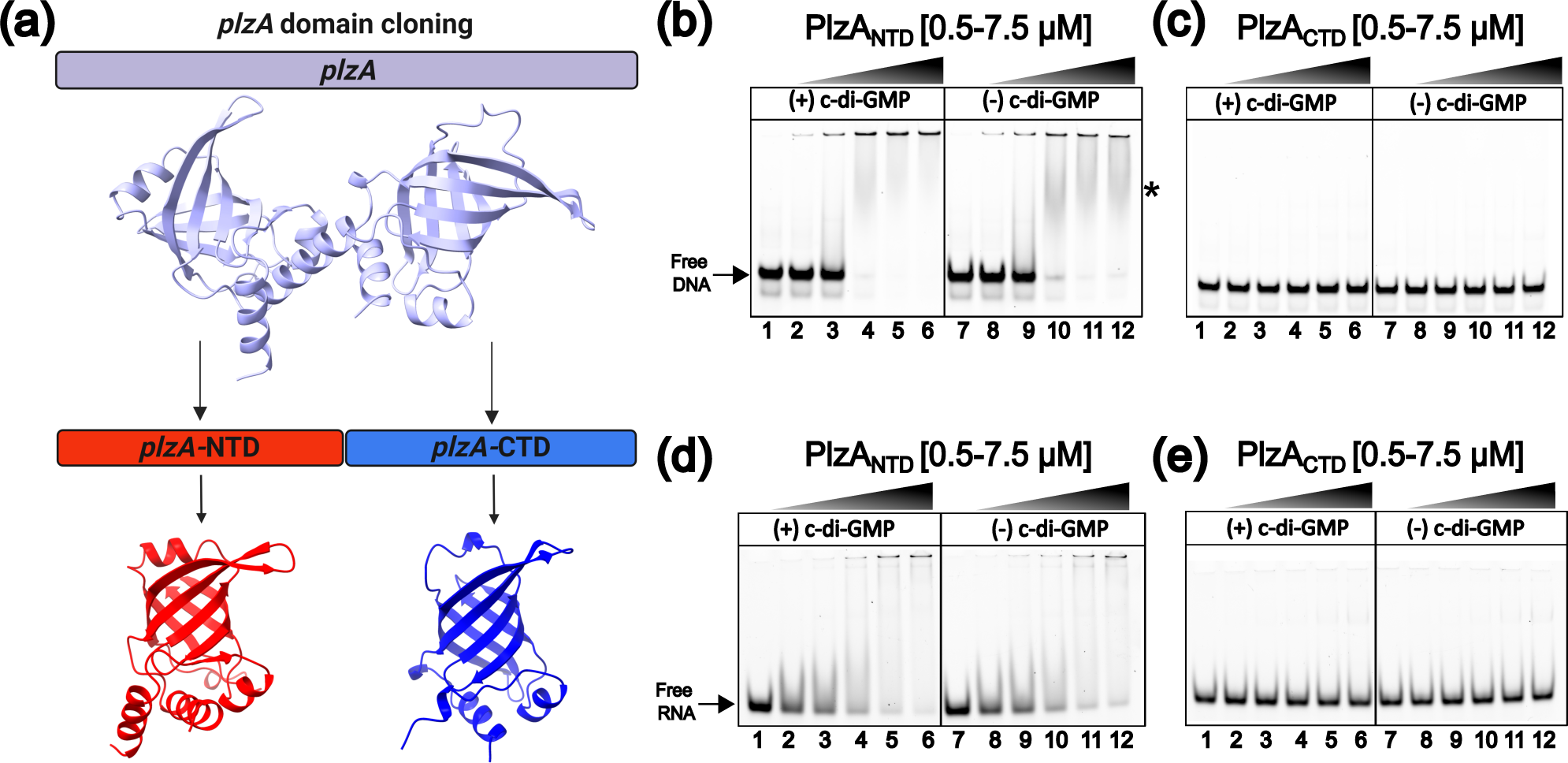
The N-terminal domain of PlzA is the nucleic acid binding domain. (**a**) The cloning strategy used to produce recombinant N (PlzA_NTD_) and C (PlzA_CTD_) terminal PlzA domains. The *plzA* gene portions corresponding to the N-terminus, *plzA*-NTD, and the C-terminus, *plzA*-CTD, were generated and cloned into an expression vector to produce recombinant PlzA_NTD_ and PlzA_CTD_ proteins. Gradient EMSAs were performed with increasing protein concentrations of PlzA_NTD_ (**b, d**) or PlzA_CTD_ (**c, e**) with (lanes 1-6) or without (lanes 7-12) 100 µM c-di-GMP and 10 nM fluorescently labeled *glpFKD*(−7/35+) dsDNA (**b, c**) or *glpFKD*(UTR) RNA (**d, e**) probe. Lanes 1 and 7 are probe-only controls. Protein concentrations were as follows: Lanes 2 and 8-0.5 µM, Lanes 3 and 9-1 µM, Lanes 4 and 10-2.5 µM, Lanes 5 and 11-5 µM, Lanes 6 and 12-7.5 µM. SUPERase•In RNase inhibitor was supplemented to a final concentration of 2 U/µL in RNA EMSA mixtures. The asterisk indicates PlzA_NTD_-dsDNA shifts. Representative gels are shown from triplicate (PlzA_NTD_) or duplicate (PlzA_CTD_) EMSAs. N= 2 or 3 independent experiments.

EMSAs were performed with labeled *glpFKD*(−7/+35) DNA probe and increasing concentrations of PlzA_NTD_ or PlzA_CTD,_ with or without 100 µM c-di-GMP. PlzA_NTD_ bound DNA independently of c-di-GMP, as supplementation with c-di-GMP did not affect binding (Figure 10B). PlzA_CTD_ contains the motifs required for c-di-GMP binding, but the PlzA_CTD_ domain did not bind to DNA with or without added c-di-GMP (Figure 10C). It was also determined that PlzA_NTD_ could bind *glpFKD*(UTR) RNA with or without c-di-GMP, while PlzA_CTD_ does not (Figure 10D and E respectively).

We also performed the LC/MS-MS analysis on PlzA_NTD_ and PlzA_CTD_ recombinant proteins to determine if any di-nucleotides were present (Figure 11A). Co-purified c-di-GMP was detected in PlzA_CTD_ but not PlzA_NTD_. The concentration of c-di-GMP present in recombinant PlzA_CTD_ was determined as 704.29 nM. A total of 66.7 µM of protein was sent for analysis indicating approximately 1.1 % of PlzA_CTD_ was bound with c-di-GMP. This indicates that PlzA_CTD_ retained the ability to bind c-di-GMP but cannot bind nucleic acid.

**FIGURE 11.**
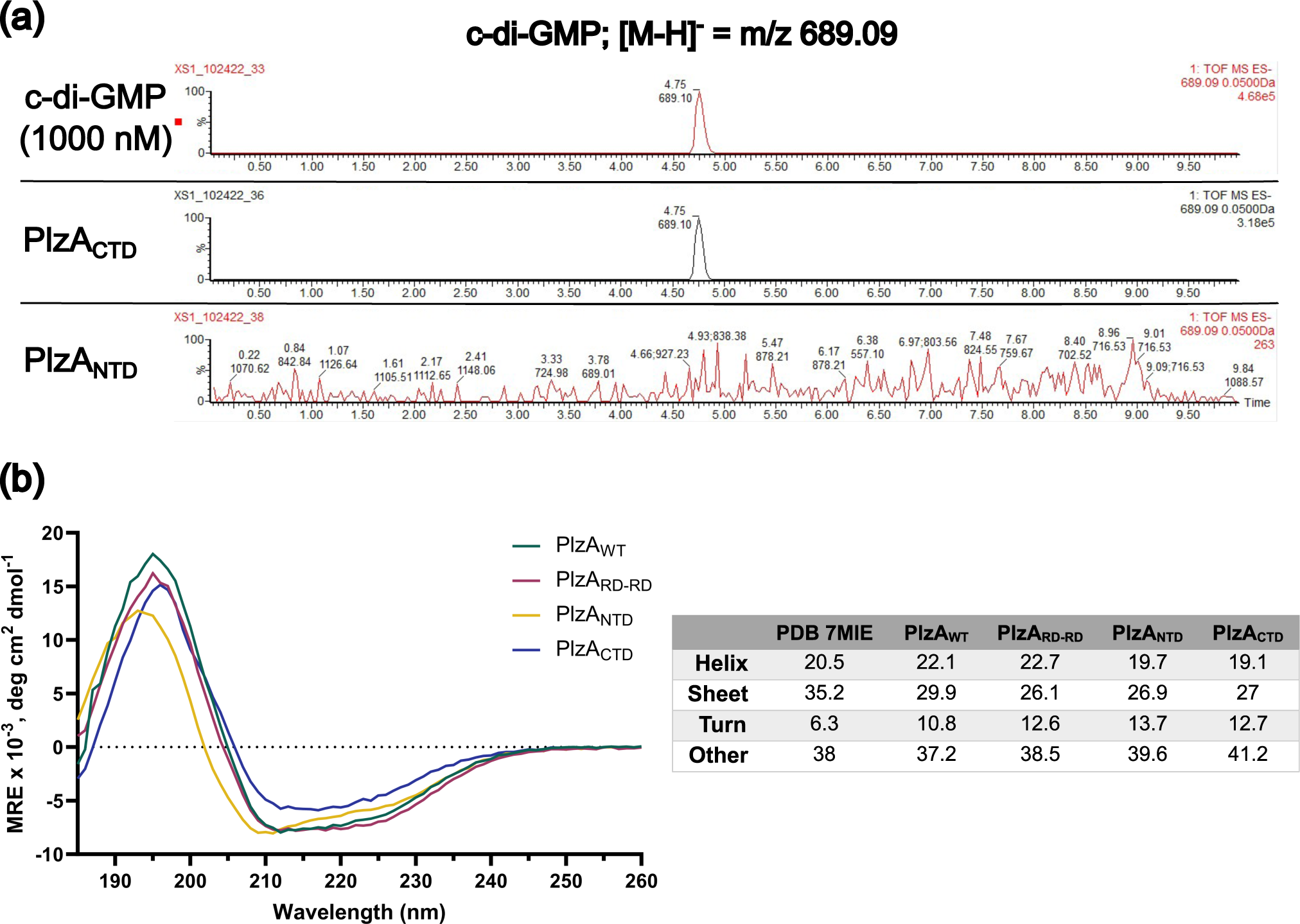
Detection of c-di-GMP in recombinant PlzA_NTD_ and PlzA_CTD_ by LC-MS/MS and circular dichroism of proteins. (**a**) Recombinant PlzA_NTD_ and PlzA_CTD_ were sent off for mass spectrometry to detect the presence of c-di-GMP. Extracted ion chromatograms of the LC-MS/MS analysis are displayed. Chromatograms depict the control c-di-GMP standard at 1 µM, recombinant PlzA_CTD_, and recombinant PlzA_NTD_. The LC-MS/MS analysis revealed that PlzA_CTD_ co-purified with c-di-GMP, but PlzA_NTD_ did not. The concentration of c-di-GMP in recombinant PlzA_CTD_ was determined from a standard curve of c-di-GMP. PlzA_CTD_ co-purified with 704.29 nM of c-di-GMP and approximately 1.1 % of the protein was bound with c-di-GMP. The c-di-GMP standard chromatogram shown is the same as in Figure 3. (**b**) Circular dichroism spectra of recombinant PlzA_WT_, PlzA_RD-RD_, PlzA_NTD_, and PlzA_CTD_ (see methods), and a summary table of the secondary structure analysis via BeStSel of recombinant proteins versus the PlzA PDB file 7MIE. CD spectra were derived from averaging the CD results from two different recombinant protein preparations for each protein tested. A buffer only control was performed and subtracted from the averaged measured recombinant protein CD spectra.

To confirm that the truncated proteins were stably folded, circular dichroism (CD) was performed, and the CD spectra were obtained for PlzA_WT_, PlzA_RD-RD_, PlzA_NTD_, and PlzA_CTD_ (Figure 11B). The CD results show that the mutated/truncated proteins retained stable structures matching that of PlzA_WT_. Therefore, the lack of nucleic acid binding by PlzA_RD-RD_ and PlzA_CTD_ was not due to improper folding. Secondary structure analysis was performed via the BeStSel server to further compare the recombinant proteins to the crystallized PlzA structure (Micsonai *et al*., 2015; Micsonai *et al*., 2018; Micsonai *et al*., 2022). All tested recombinant proteins show some differences from the calculated secondary structures of the 7MIE PDB file. These deviations are primarily in the percentage of beta-sheet and turn secondary structure present. We note that the His-Tag was not cleaved prior to CD analysis of the recombinant proteins, and the resultant deviations could be due to the tag. The tag was considered in all calculations for CD. Overall, we can deduce that protein folding is not the cause of the lack of binding by PlzA_RD-RD_ or PlzA_CTD_.

### Computational docking of PlzA-nucleic acid interactions

To gain a better understanding of the interactions between PlzA and DNA or RNA, we performed a computational docking analysis with the HDOCK server (Huang and Zou, 2014; Yan *et al*., 2017a; Yan *et al*., 2017b; Yan *et al*., 2020). The *glpFKD*(−7/+35) DNA and *glpFKD*(UTR) RNA sequences were input as ligands, while the PlzA crystal structure (PDB ID: 7mie) was input as the receptor molecule. The modeling results indicate that interactions between PlzA and *glpFKD*(−7/+35) DNA are facilitated primarily through the N-terminal domain (Figure 12A). In the representative docking data, beta-strands six and seven of the N-terminus are extended and protrude outward, intercalating a major groove of the DNA 5’ to the protein docking site (Figure 12A top panel). Polar and positively charged amino acids, including N110 and K111, are located at the loop region where beta-strands 6 and 7 converge, with the potential to directly interact with bases of the DNA (Rooman *et al*., 2002; Sathyapriya and Vishveshwara, 2004). Interactions with a minor groove of the DNA are predicted to occur through the loop region of beta-strands 2 and 3 and an unstructured linker region (Figure 12a bottom panel). Several positively charged residues, including arginine (R32 and R38) and lysine (K81), as well as an aromatic phenylalanine residue (F45), are predicted to directly interact with the minor groove and backbone of the DNA. An additional lysine residue K87, which is in the unstructured region, protrudes and interacts directly with bases of another major groove of *glpFKD*(−7/+35). No residues of the C-terminal domain are predicted to interact directly with the DNA; however, four residues are interfacing the DNA (K161, I213, D258, and N261). Residue K161 hovers over the DNA backbone potentially interacting nonspecifically via electrostatic interactions. We note that the docking analysis produced several potential models that differed in the location of where residues interact with either the minor or major groove. The model presented is the highest-ranking model. Overall, the computational docking data supports our EMSA data showing that PlzA_NTD_ binds DNA while PlzA_CTD_ does not (Figures 10B and C) and that PlzA interacts predominately with the major groove of DNA (Figure 5).

**FIGURE 12.**
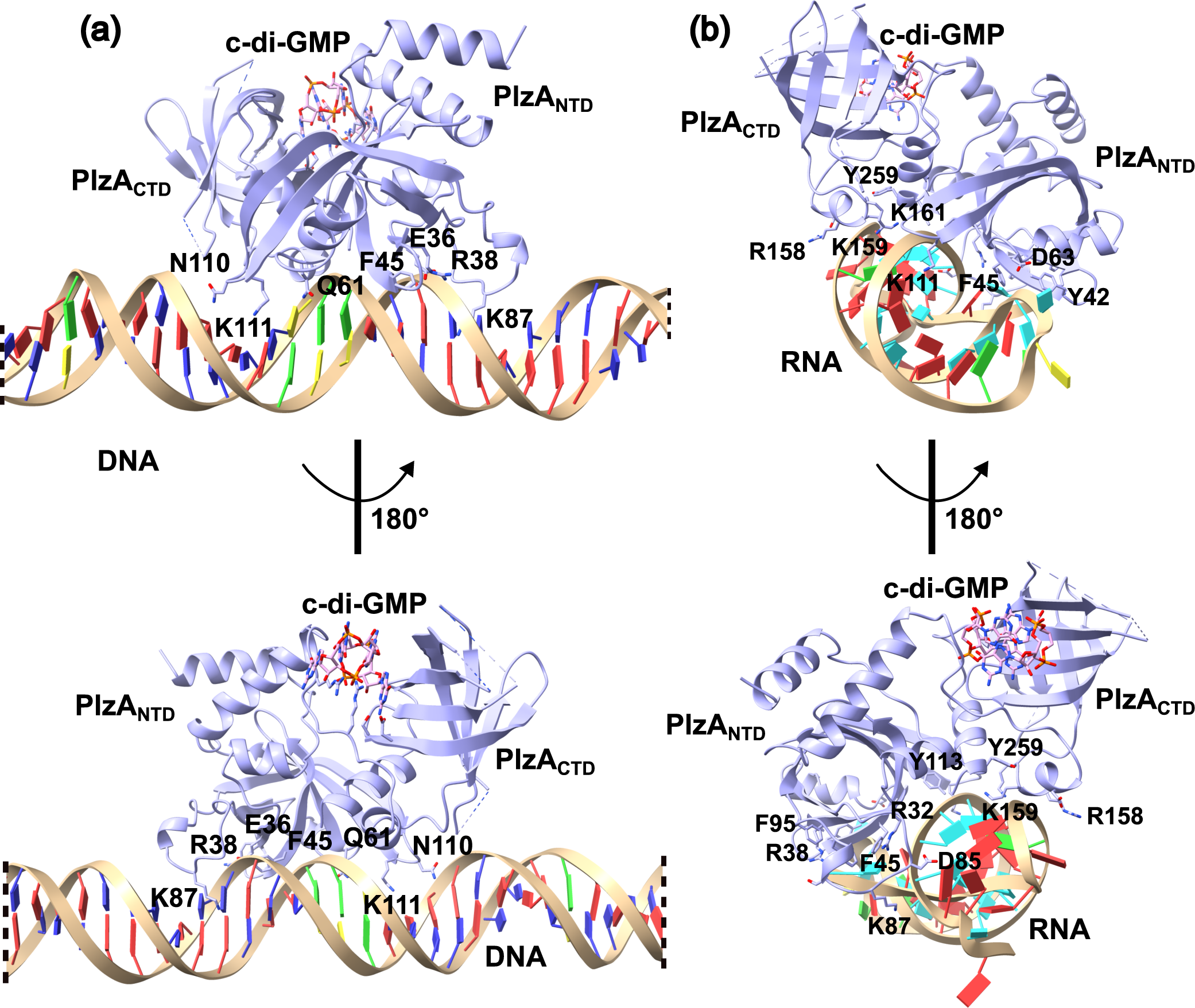
Computational docking analysis of PlzA with *glp* DNA and RNA. Computational docking was performed using the HDOCK server with PlzA and either (**a**) the 42 bp *glpFKD*(−7/+35) dsDNA or (**b**) the *glpFKD*(UTR) RNA sequence. In (**a**) only 26 of 42 bp of the *glpFKD*(−7/+35) sequence is shown. The dashed lines represent where bases were left out of the figure to zoom in on the protein-DNA interactions. The dsDNA model was linear, and removal of the bases has no impact on the interpretation of the data. The top predicted model from each computation is shown. The HDOCK docking and confidence scores for each model were −210.60 and 0.77 for PlzA-*glp* DNA, and −280.71 and 0.93 for PlzA-*glp* RNA, respectively. The protein is rotated 180° from the top to the bottom panels. Select amino acid residues that interface the nucleic acids are labeled.

The docking analysis was also performed with PlzA_WT_ and *glpFKD*(UTR) RNA (Figure 12B). Again, the N-terminal domain of PlzA facilitates most of the interactions. Similar regions of the protein that interacted with dsDNA are implicated in the interaction with RNA including the residues of the loop region of beta-strands 6 and 7 (K111 and Y113), the loop region of beta-strands 2 and 3 (R38, Y42, and F45) and the unstructured linker region (R32, D85, K87, F92, and F95). Unlike the docking model with dsDNA, the PlzA-RNA docking model indicates more involvement of the C-terminal domain in RNA binding. Several residues of alpha-helixes 5 (D155, R158, K159, and K161) and 6 (D258 and Y259) in the C-terminal domain of PlzA, are predicted to interact with RNA. Although our EMSA results show that PlzA_CTD_ does not bind any tested nucleic acids, the docking results indicate it could help stabilize PlzA_NTD_-nucleic acid interactions or could be involved in target recognition. Guided by the docking data, we are in the process of generating site-direct PlzA mutant proteins that will explore residues predicted to be involved in nucleic-acid binding.

Computational docking was attempted with PlzA and ss*glpF*, but the HDOCK server could not complete the analysis. Given that PlzA bound dsDNA and RNA better than ssDNA, it is interesting to note that the computational modeling predicts the RNA to fold into a double stranded secondary structure. Combined with our EMSA data, this suggests that PlzA could have a preference towards double stranded nucleic acids and/or secondary structures.

## DISCUSSION

Cyclic-di-GMP has wide-ranging effects on many bacterial processes. The responses to c-di-GMP are commonly mediated through c-di-GMP binding effector proteins. PlzA is the only chromosomally and universally encoded c-di-GMP binding protein of the Lyme disease spirochetes. It has been shown to be crucial for borrelial infection processes and in the regulation of the metabolism of alternative carbon sources during colonization of vector ticks (Pappas *et al*., 2011; Pitzer *et al*., 2011; He *et al*., 2014; Mallory *et al*., 2016; Zhang *et al*., 2018; Groshong *et al*., 2021). Despite its singularity and importance to *B. burgdorferi*, little was previously known about the biochemical functions of PlzA outside of binding c-di-GMP. Here, we provide evidence that PlzA is a novel c-di-GMP-dependent nucleic acid-binding protein. While PlzA lacks an obvious canonical DNA binding motif, we hypothesized that PlzA might be a nucleic acid binding protein given its role in the regulation of the *glpFKD* operon. Binding was only observed by the wild-type PlzA protein but not the mutant, PlzA_RD-RD_, which is incapable of binding c-di-GMP. It has previously been shown that structural rearrangements are induced upon the binding of c-di-GMP to PlzA (Mallory *et al*., 2016; Groshong *et al*., 2021). Without bound c-di-GMP, The N- and C-terminal domains of PlzA can flex on an unstructured linker, potentially inhibiting the stable interactions that are necessary for DNA and RNA binding. This hypothesis is supported by the fact that crystallization of PlzA was only achieved when saturated with c-di-GMP (Singh *et al*., 2021). We surmise that binding of c-di-GMP to the C-terminal PilZ domain results in a change of PlzA into a more rigid structure, which permits nucleic acid binding through residues of the N-terminal domain.

We identified the N-terminal domain of PlzA as the nucleic acid binding domain and that PlzA_NTD_ bound DNA independently of c-di-GMP. In contrast, the C-terminal domain, PlzA_CTD_, did not bind any tested nucleic acids, although it still bound c-di-GMP. While PlzA_NTD_ can interact with nucleic acids, these interactions were not as strong as PlzA_WT_. Binding of nucleic acid by PlzA_NTD_ resulted in smeared shifts that lacked defined bands. This suggests instability of the PlzA_NTD_-nucleic acid complexes resulting in dissociation of the complexes during electrophoresis. Noting that binding of c-di-GMP appears to stiffen interactions between the N- and C-terminal domains, we hypothesize the C-terminal domain may stabilize the interactions of PlzA_NTD_ with nucleic acids when c-di-GMP is bound. On the other hand, we posit that the flexibility of apo-PlzA results in the C-terminal domain disrupting the ability of the N-terminal domain to stably bind DNA. Although the N- and C-terminal PlzA domains are similar, structural comparisons of PlzA_NTD_ and PlzA_CTD_ reveal distinct differences (Figure 10A). The N-terminus of PlzA_NTD_ contains an alpha-helix that is absent in the N-terminus of PlzA_CTD_. Additionally, an alpha-helix is present at the beta-barrel pore of PlzA_NTD_ but not in the PlzA_CTD_. These differences suggest distinct roles in PlzA-nucleic acid binding dynamics. We are currently exploring the sequence motifs recognized by PlzA and the exact amino acid residues involved in nucleic acid binding.

Previous studies found that PlzA is a monomer in solution regardless of c-di-GMP (Mallory *et al*., 2016). While many DNA-binding proteins bind DNA as dimers, alternative binding models exist (Kim and Little, 1992; Kohler *et al*., 1999). It has been shown that some proteins are capable of binding sequentially as monomers which then multimerize on the DNA via protein-protein interactions, such as some members of the Leucine zipper and helix-loop-helix zipper families, the LexA repressor of *E. coli*, and BpaB of *B. burgdorferi* (Kim and Little, 1992; Kohler *et al*., 1999; Burns *et al*., 2010). Our studies revealed that increasing c-di-GMP and PlzA_WT_ concentrations resulted in progressively slower migration of PlzA-*glpFKD*(−7/+35) DNA complexes. The decreases in the migration of the complexes that are observed with increasing protein concentration may indicate a stoichiometry that is not 1:1, or the observed smearing may be the result of dynamic equilibrium of the complexes in the gels during electrophoresis (Fried and Crothers, 1981; Fried, 1989). While increasing c-di-GMP enhances PlzA binding, there could be a threshold where too much c-di-GMP is inhibitory and causes dissociation of PlzA from nucleic acids. A dimerization or protein-protein interaction residue/motif/domain remains to be identified in PlzA, and work is ongoing to address the DNA-binding mode and stoichiometry of PlzA-nucleic acid complexes.

We also observed probe signal in the gel wells at high protein and c-di-GMP concentrations. This could be due to protein aggregation. Rather than simply being a negative consequence of environmental stress, recent studies indicate a role for protein aggregation in regulation. Specifically, certain conditions can induce a protein to aggregate, and thus be sequestered, altering gene expression to permit bacterial adaptation to a specific environment (Schramm *et al*., 2019). Whether high levels of c-di-GMP promote such processes remains to be tested, but c-di-GMP levels certainly alter PlzA binding affinity.

Our study sought to identify if PlzA can bind nucleic acids and what role c-di-GMP plays in that function. The data are conclusive that PlzA indeed binds DNA, both double and single-stranded, and RNA, with a preference for dsDNA and RNA. PlzA also displayed higher affinity for certain nucleic acid types over others, and PlzA-*glp* DNA binding was competed away by specific competitors but not poly-dI-dC. Further, the nucleic acid binding at the *glpFKD* locus by PlzA is dependent on c-di-GMP. A caveat of the data is that the calculated dissociation constants for PlzA-nucleic acids are lower than typically observed for protein-nucleic acid interactions. The higher dissociation constants could be attributed to our method of choice, EMSA, as some potential dissociation and protein aggregation were observed. Attempts were made with various alterations aimed to improve the stability of the complexes during electrophoresis, such as altering the gel percentage, pH of the running buffer, salt/ion concentrations of the reaction buffer, and addition of detergents to the reaction (Hellman and Fried, 2007). Altering the pH of the running buffer to 8.0 decreased the aggregation in the wells but smearing still occurred at higher protein and c-di-GMP concentrations (data not shown). The experimental conditions presented and reported here yielded the most consistent results.

PlzA bound to a sequence adjacent to the promoter and extending 35 bp into the 5’ UTR of *glpF*. This site has been shown to positively affect *glpFKD* transcript levels and is a site where two other borrelial regulators, SpoVG and BadR, also bind (Grove *et al*., 2017; Savage *et al*., 2018; Saylor *et al*., 2023; Zhang *et al*., 2024). BadR, the *<u>B</u>orrelia* Host <u>Ad</u>aption <u>R</u>egulator, has recently been identified as a repressor of *glpFKD*, while PlzA has previously been shown to exert both positive and negative effects on *glpFKD*, at high and low levels of c-di-GMP respectively (Zhang *et al*., 2018; Zhang *et al*., 2024). Although SpoVG can bind the DNA and RNA of *glpFKD*, the transcriptional consequences of these interactions are unknown (Savage *et al*., 2018; Saylor *et al*., 2023). How these proteins work in tandem to regulate glycerol catabolism is currently being investigated. Multi-layered regulation of glycerol catabolism is found in other bacteria, indicating this is not a phenomenon unique to *Borrelia* (Bong *et al*., 2019).

Our data show that PlzA is a DNA and RNA binding protein that is c-di-GMP dependent. Obvious questions that remain are what sequence motif(s) PlzA recognizes and what additional binding sites may exist in the *B. burgdorferi* genome. Our studies were unable to identify a sequence motif in the *glpFKD* intergenic region but were able to identify that PlzA interacts mainly with the major groove of DNA. We also saw a preference of PlzA for longer sequences and suggest PlzA may recognize gene targets in a sequence independent manner. The lesser affinity for shorter sequences could indicate that PlzA might recognize structured nucleic acids, as shorter DNA is less likely to form structured conformations. Given that poly-dI-dC did not abolish binding, PlzA likely exhibits preferential affinity to certain sequences. Whether the preferential affinity is due to a sequence motif or structural motifs, remains to be determined. What is clear, from multiple studies, is that the cellular effects exerted by PlzA are specific.

A logical step to address these questions is to perform chromatin immunoprecipitation with sequencing (ChIP-Seq). Given that PlzA binding is reliant on an effector molecule, c-di-GMP, there are some caveats in performing this assay at this stage. First, the activating stimulus for the Hk1/Rrp1 pathway is currently unknown. As a result, it is unknown how levels of c-di-GMP fluctuate in culture medium and across growth phases, and therefore how much c-di-GMP is produced in a bacterial cell at a given time. This can affect the ability to obtain consistent ChIP results. The tendency of protein to occupy each site will likely vary with culture conditions. Therefore, optimal experimental conditions must be determined before obtaining and drawing conclusions about PlzA binding sites derived from ChIP experiments.

Next, as mentioned above, two other proteins have been found to bind the same site within *glpFKD*, SpoVG and BadR(Savage *et al*., 2018; Saylor *et al*., 2023; Zhang *et al*., 2024). Competition between these proteins could also impact the ChIP data. Additionally, the molecular consequence of nucleic acid binding remains to be determined. We are actively working to address these questions by characterizing several PlzA site-direct mutant proteins incapable of binding nucleic acids to determine impacts on *B. burgdorferi* physiology and gene regulation. Such work will also aid in identifying targets of PlzA regulation to better guide the identification of additional binding sites.

During the preparation of this manuscript, Van Gundy et al. published a report identifying RNA chaperone activity by PlzA (Van Gundy *et al*., 2023). The authors concluded that this activity is independent of c-di-GMP, which deviates from our results showing that c-di-GMP is required for nucleic acid binding. Those authors acknowledged this and suggested that differences in methodologies and conditions might have contributed to the differences in observed results. Some of these differences include using non-borrelial derived RNAs, a different PlzA site-mutant, equimolar concentrations of c-di-GMP, and filter binding assays used to calculate dissociation constants.

We chose to evaluate saturating c-di-GMP concentrations to evaluate the maximal nucleic acid binding activity of holo-PlzA and mitigate potential variability in the calculated apparent K_D_ values. Van Gundy et al. did not assess whether their PlzA protein preparations co-purified with c-di-GMP, which could have contributed to their observed binding by apo-PlzA. Unlike their study, we did not observe binding by the PlzA_RD-RD_ mutant, which cannot bind c-di-GMP, under any tested condition. Given the stabilizing effect c-di-GMP binding has on the structure of PlzA (only the holo form was able to be crystallized), the ability of their mutant and apo-PlzA to bind RNA merits further mechanistic investigation. Lastly, the goals for our studies differed greatly as they sought to study RNA chaperone activity, while we set out to investigate the ability of PlzA to bind nucleic acids. Our two studies, however, corroborate that PlzA can bind RNA and suggest that this function plays a role in regulating borrelial gene expression. We agree that further investigation is required to unravel the molecular mechanisms of PlzA interactions with nucleic acids, and the roles that those interactions play in gene regulation in this important pathogen.

It is known that the affinity and specificity of proteins for target nucleic acids are impacted by co-effectors (Carey, 2022). We observed that adding exogenous c-di-GMP increased binding affinity of PlzA for *glp*-derived DNA and RNA. The localization of c-di-GMP and associated proteins in *B. burgdorferi* is currently unknown. Whether the c-di-GMP signaling network is global or localized will be key to understanding how c-di-GMP induces PlzA effector functions. This is actively being investigated by our laboratory. Studies in other bacteria with multiple PilZ domain proteins have suggested that the differential affinity of the various PilZ receptors for c-di-GMP sequentially regulates the processes that respond to c-di-GMP (Pultz *et al*., 2012; Junkermeier and Hengge, 2023). Given that PlzA is the only PilZ domain protein in most *B. burgdorferi* isolates, an alternative to this could be that cellular levels of c-di-GMP alter the affinity and/or specificity of PlzA.

We propose a model in which c-di-GMP availability alters the functionality of PlzA, and thus, the targets regulated by PlzA could depend on the c-di-GMP levels available at different stages of the enzootic life cycle (Figure 13). During vertebrate infection, no c-di-GMP is produced, favoring apo-PlzA functions such as RNA chaperone activity (Van Gundy *et al*., 2023). When c-di-GMP is present, holo-PlzA can also anneal RNAs and potentially interact with RNAP (Grassmann *et al*., 2023; Van Gundy *et al*., 2023). At lower levels of c-di-GMP, PlzA nucleic-acid binding affinity is low for target loci. As c-di-GMP levels increase further, binding affinity for target nucleic acids is increased and potential multimerization of PlzA molecules on DNA could occur.

**FIGURE 13.**
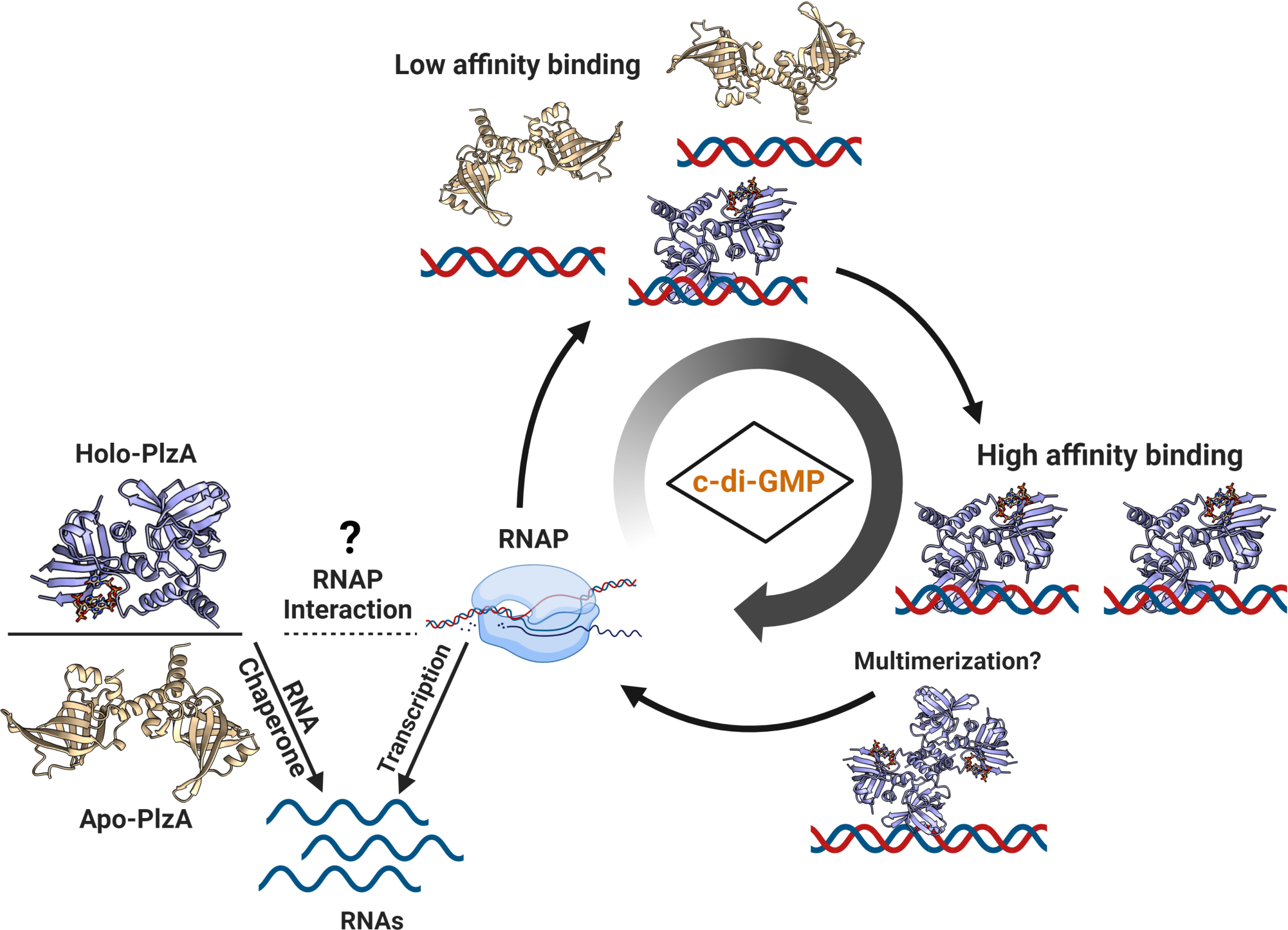
Model depicting the modulation of PlzA function by levels of c-di-GMP. Apo and holo-PlzA are hypothesized to have distinct functions in the vertebrate and tick hosts respectively. Recent studies identified PlzA RNA chaperone activity that is c-di-GMP independent (Van Gundy *et al*., 2023). In environments with no c-di-GMP, such as the vertebrate host, apo-PlzA engages in RNA chaperone activity to modulate gene expression required for survival in the vertebrate host. The phosphodiesterases PdeA and PdeB are likely to be highly active upon transmission and into vertebrate infection, lowering c-di-GMP levels and facilitating apo-PlzA functions (Sultan *et al*., 2010; Pitzer *et al*., 2011; Sultan *et al*., 2011; Novak *et al*., 2014; Groshong *et al*., 2021). Phosphodiesterase activity is likely to occur at all stages to regulate c-di-GMP levels and indirectly, PlzA function. When Hk1 is activated (signal unknown), Rrp1 is phosphorylated and produces c-di-GMP. Holo-PlzA can also engage in RNA chaperone activity by annealing RNAs, and potentially interact with RNAP (Grassmann *et al*., 2023; Van Gundy *et al*., 2023). We propose that levels of c-di-GMP impact the specific genes regulated by PlzA through altering nucleic acid binding affinity. At lower c-di-GMP levels, few holo-PlzA monomers exist resulting in weak PlzA binding affinity for certain target loci. As levels of c-di-GMP increase, more holo-PlzA molecules bind target genes with higher affinity, and PlzA could potentially multimerize on DNA.

In conclusion, PlzA is a c-di-GMP-dependent nucleic acid-binding protein. While other c-di-GMP binding receptors have been identified as DNA-binding proteins, PlzA is the first that has been demonstrated to also bind RNA, thus expanding the functional diversity of c-di-GMP binding proteins. Given that PlzA can bind to both DNA and RNA, it could have a role in coordinating transcriptional complexes, and it supports the notion that PlzA can interact with RNAP (Grassmann *et al*., 2023). Our study examined PlzA nucleic acid binding at a locus known to be affected by PlzA, the *glpFKD* catabolism operon, but sites of higher affinity could exist. Our work identifying PlzA as a novel nucleic acid-binding protein provides a basis for its functional mechanism. This will further inform our understanding of how *B. burgdorferi* regulates gene expression in response to environmental cues and help unravel the PlzA regulon.

## MATERIAL AND METHODS

### Routine manipulations and molecular cloning

Oligonucleotides used in this study are listed in Table 2 and were ordered from Integrated DNA Technologies (IDT). *B. burgdorferi* genomic DNA was purified utilizing the E.Z.N.A Tissue DNA kit (OMEGA). Standard and high-fidelity PCR were performed with 2X DreamTaq Green PCR Master Mix (Thermo Scientific) or Q5 High-Fidelity 2X Master Mix (NEB), respectively. Plasmids were isolated using QIAprep Spin Miniprep kits (Qiagen) according to the Miraprep protocol (Pronobis *et al*., 2016). All plasmid constructs were transformed into chemically competent *E. coli* Top10 (Invitrogen) or DH5α (Invitrogen) strains for cloning and plasmid maintenance. Positive clones were identified by colony PCR screening with the appropriate primer sets, and DNA sequencing was performed by Eurofins Genomics LLC (Louisville, KY).

**Table 2.**
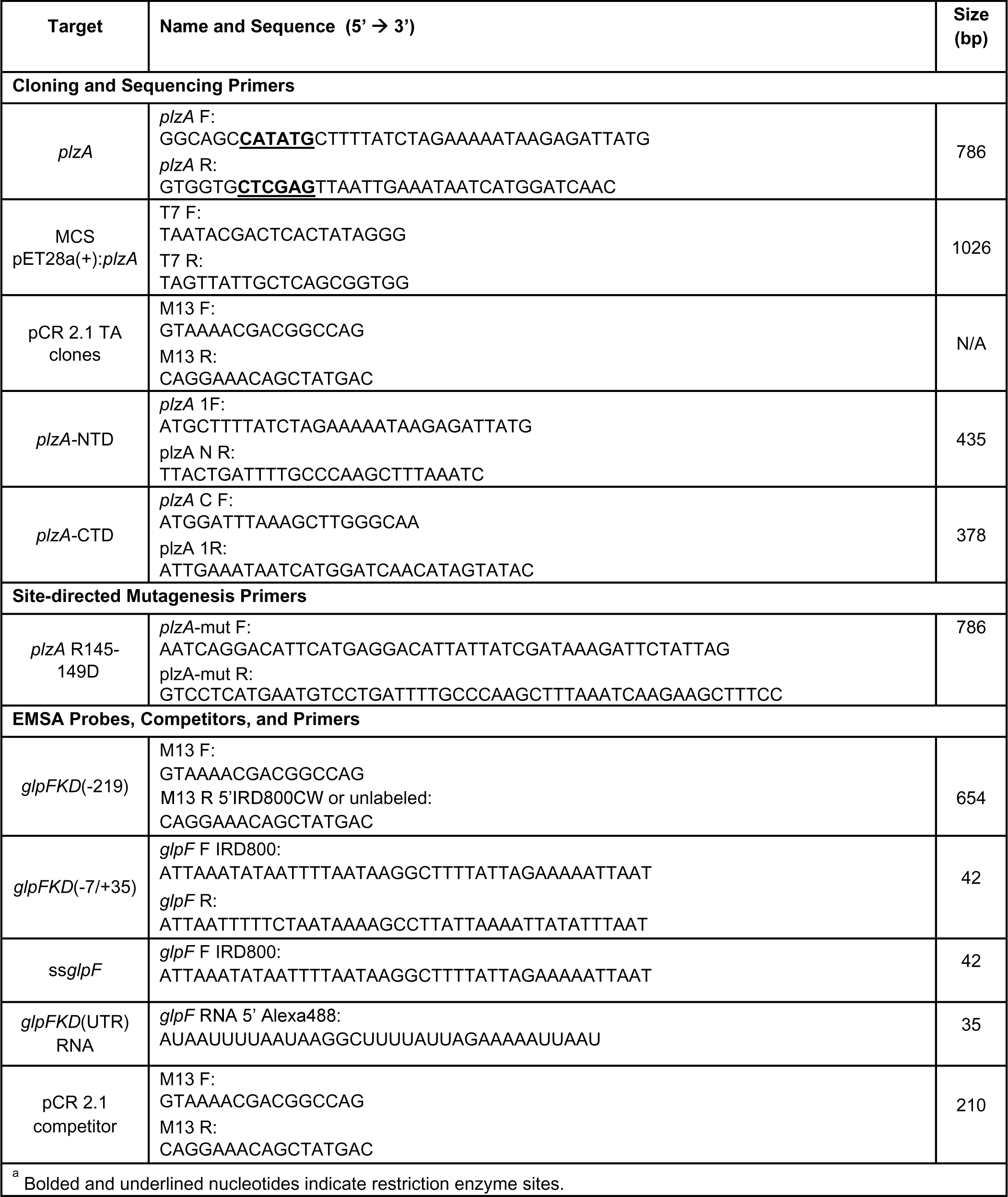
Oligonucleotides used in this study.

### Plasmid construction

The wild-type (WT) *plzA* gene from *B. burgdorferi* strain B31, was cloned between the Ndel/XhoI sites of pET28a(+) and thus fused in-frame with an amino-terminal 6xHis-tag. Truncated *plzA* genes, named *plzA*-NTD and *plzA-*CTD encoding the N-terminal (residues 1-141) and C-terminal (residues 142-261) domains of PlzA, respectively, were generated via gene synthesis and cloned to produce 6xHis-tagged proteins by Genscript (Piscataway, NJ). Briefly, the synthesized truncations were cloned into the pET28b(+) vector with *plzA*-NTD cloned between the BamHI/NotI sites producing an amino-terminal 6xHis-tag fusion, while *plzA* CTD was cloned between the NcoI/XhoI sites producing a carboxy-terminal 6xHis-tag fusion.

### Generation of the site-directed PlzA R145D-R149D mutant

Site-directed mutagenesis by overlap extension PCR was used to generate a PlzA mutant that cannot bind c-di-GMP. Two-point mutations were introduced into a pTrcHis TOPO (Invitrogen) construct containing the *plzA* gene. The successful introduction of the mutations was confirmed by DNA sequencing. The site-directed PlzA double mutant is denoted as PlzA_RD-RD_. The oligonucleotide primers used for site-directed mutagenesis are listed in Table 2.

### Recombinant protein overexpression and purification

For recombinant protein expression, *E. coli* Rosetta 2(DE3)(pLysS) (Novagen) chemically competent cells were used for PlzA_WT_, PlzA_RD-RD_, and N-terminal PlzA domain (PlzA_NTD_), while One Shot BL21 Star (DE3) cells were used for the C-terminal PlzA domain (PlzA_CTD_). Overnight bacterial cultures were diluted 1:100 into Super broth (tryptone 32 g, yeast extract 20 g, NaCl 5 g per liter) supplemented with the appropriate antibiotics (kanamycin-50 μg/ml, carbenicillin-100 μg/ml, and/or chloramphenicol-30 μg/ml). The cultures were allowed to grow to an OD_600_ of 0.5-1.0 and protein expression was subsequently induced by the addition of 0.25 mM (PlzA_NTD_ only) or 0.5 mM isopropyl-β-D-thiogalactopyranoside (IPTG). Induced cultures were incubated at either 37°C for 3-4 h (PlzA_WT_ and PlzA_CTD_), 29°C for 4-6 h (PlzA_WT_), or overnight at room temperature (PlzA_RD-RD_ and PlzA_NTD_). The cells were harvested by centrifugation at 5400 x *g* for 30 min at 4°C, and the cell pellets were frozen at −80°C until protein purification.

For protein extraction, cell pellets were thawed on ice and resuspended in equilibration buffer (20 mM sodium phosphate, 300 mM NaCl, 10 mM imidazole, pH 8.0) and lysed via sonication. Lysates were clarified by centrifugation at 23700 x *g* for 20 min at 4°C, and the supernatant was retained. When further clarification was required due to lysate viscosity, the supernatants were passed through a sterile 0.22 μM Millex-GS Syringe Filter Unit (MilliporeSigma). Protein purification was performed via column affinity chromatography using either HisPur Ni-NTA resin (Thermo Scientific) or the Magne-His purification system (Promega Corporation) for PlzA_RD-RD_ per manufacturer protocol. Eluted protein solutions were dialyzed overnight into either EMSA buffer (50 mM Tris-HCl [pH 7.5], 50 mM KCl, 1 mM dithiothreitol (DTT), 1 mM ethylenediaminetetraacetic acid (EDTA) [pH 8.0], 1 mM phenylmethanesulfonyl fluoride (PMSF), 10% glycerol (v/v), and 0.01% Tween-20) or 20 mM sodium phosphate, pH 7.2. The dialyzed proteins were then concentrated using Amicon Ultra 3K (PlzA_N_ and PlzA_C_) or 10K (PlzA_WT_ and PlzA_RD-RD_) centrifugal filter units (MilliporeSigma). Aliquots of dialyzed proteins were assessed for purity by SDS-PAGE and Coomassie brilliant blue staining. Final recombinant protein concentrations were determined using Quick Start Bradford Protein Assays (Bio-Rad) with bovine serum albumin as the reference protein for standard curves. Purified protein aliquots were stored at −80°C.

### Nucleic acid fragments design and generation

All probe sequences were derived from the genomic sequence of *B. burgdorferi* type strain B31, and the corresponding primers are reported in Table 2 (Fraser *et al*., 1997; Casjens *et al*., 2000). Large and small DNA substrates were used in this study. Larger fluorescently tagged fragments >60 bp were generated by PCR amplification of the target sequences from B31 genomic DNA and subsequent cloning into the pCR 2.1 TA vector (Invitrogen) following TOPO TA protocols. Subsequently, target-specific or M13 F and R IRDye 800-labeled primers were used for PCR of the plasmid templates to produce labeled DNA fragments for binding studies. All PCR-generated probes were treated with exonuclease I (NEB) according to the manufacturer’s protocol to remove single-stranded DNA, followed by ethanol precipitation. The DNA was resuspended in molecular-grade water (Ambion) and quantified by Nanodrop UV spectrometry.

Smaller probes <60 bp long were produced by annealing complementary unlabeled and IRDye 800-labeled oligonucleotides at equimolar concentrations at 95°C for 5 min and were gradually cooled overnight at RT. The same annealing procedure was done for unlabeled competitors. All probes and competitors were aliquoted and stored at −20°C until further use. Unlabeled and 5’ IRDye 800 (LI-COR Biosciences) labeled oligonucleotides were synthesized by IDT. Alexa Fluor 488 conjugated RNA probes corresponding to the transcribed regions of the respective DNA probes were also synthesized by IDT.

### Electrophoretic mobility shift assays (EMSA)

EMSA reactions were performed in EMSA buffer at RT with a final concentration of dsDNA probe of 10 nM or RNA probe of either 10 nM or 50 nM. EMSA reaction mixtures were supplemented with 3’5’ c-di-GMP (Sigma-Aldrich) at various concentrations ranging from 0 to 100 µM. c-di-GMP was not added to the gel or running buffer. Experiments using equimolar concentrations of c-di-GMP are relative to the final protein concentration in the reaction. Proteins were co-incubated with c-di-GMP for 5 min before the addition of any nucleic acids. Prior to the addition of labeled probe, the non-specific competitor poly-dI-dC (Roche) was added to the EMSA reactions at a final concentration of 2.5 ng/µL, and the reaction mix was allowed to incubate another 5 min (Larouche *et al*., 1996). Labeled probes were added last, and the reactions were subsequently incubated for an additional 10 min prior to loading onto gels. Competition EMSAs were conducted similarly but with the unlabeled, specific competitor added after poly-dI-dC and then allowed to incubate with the reaction prior to the addition of the labeled probe. Regarding RNA EMSAs, when RNase contamination was evident in recombinant protein preparations, RiboGuard (Lucigen) or SUPERase•In (Ambion) RNase inhibitor was added to the EMSA mixtures before the addition of RNA to a final concentration of 4 U/µL and 2 U/µL respectively to inhibit RNase activity. The final reaction volume was 10 µL prior to the addition of loading dye.

For visualization during electrophoresis, 2 µL of EMSA loading dye (0.8 mg/mL Orange G, 15 mg/mL Ficoll 400) was added to each reaction prior to gel loading. Novex TBE 6 or 10% gels (Invitrogen) were pre-run in 0.5x TBE running buffer (pH ∼8.4 +/-0.5), at 100V for a minimum of 30 min. Due to the complexes of the larger *glpFKD*(−219) probe and higher concentrations of PlzA not exiting the well, the pH of the running buffer was adjusted to a pH of 8.0 immediately prior to electrophoresis which allowed the complexes to enter the gel matrix. The pH was not adjusted for EMSAs using nucleic acid fragments <60 bp as complexes readily exited the well at the standard pH of 0.5x TBE running buffer. The entire reaction mixture was then loaded onto the pre-run gels and resolved at 100 V for 60-90 min at RT. EMSA images were acquired with a ChemiDoc Imaging System (Bio-Rad).

### Densitometry and statistics

Densitometry of EMSAs was performed using Image Lab 6.1 (Bio-Rad) software. Analyses were performed on triplicate EMSAs. Lanes as well as free and shifted bands were added manually, and intensity values were reported as band percentage. For dsDNA EMSAs, free dsDNA band percentages were normalized to the probe only control band percentage. The free DNA band percentage values were then used to determine the percent shifted of the respective probe relative to the free probe. For RNA and ssDNA EMSAs, lanes and bands were selected manually, and the shifted bands were analyzed to obtain the percent shifted values. Background and any ssDNA probe percentage that was shifted were subtracted from the percent totals. For competition EMSAs, the band intensity values of each band in a lane were determined manually, and the free probe band intensity values determined. Any probe signal stuck in the gel wells, attributed to potential protein aggregation, were not considered shifts, and thus omitted from densitometric analysis.

To calculate the apparent dissociation constant (K_D_), nonlinear regression analysis was performed to determine the best-fit values for each experimental condition tested. Briefly, protein concentrations were plotted against the shifted band percentage values obtained from densitometry using the one-site specific binding setting in Prism GraphPad 9.5.1. Confidence was set at 95% with the analysis considering each replicate value from triplicate EMSAs performed for the analysis. Either a Welch’s t-test or Welch ANOVA followed by a Dunnett’s T3 multiple comparisons test, was used to compare fits.

### Major and Minor Groove Binding Assay

The IRDye 800 labeled *glpFKD*(−7/+35) DNA probe was used in the major and minor groove binding assay. The reactions were assembled similarly to the EMSA reactions described above. Briefly, 2.5 µM of WT PlzA was incubated with 100 µM c-di-GMP in EMSA buffer for 5 minutes. The *glpFKD*(−7/+35) DNA probe (10 nM) was then added to the reaction mixture which was allowed to incubate for an additional 10 minutes at room temperature. After the probe incubation, 0.25-250 µM of either methyl green (major groove binder) or actinomycin D (minor groove binder) were added to the reaction mixture and allowed to incubate for 5 minutes at room temperature. EMSA loading dye (0.8 mg/mL Orange G, 15 mg/mL Ficoll 400) was added to each reaction, and the reaction mixtures were subsequently loaded onto pre-run 10% Novex TBE gels. Gels were resolved and images acquired as described above for EMSAs.

### Docking analysis and protein modeling

Computational modeling of holo-PlzA (PDB ID: 7mie) in complex with the nucleic acids of interest was performed using the HDOCK server: http://hdock.phys.hust.edu.cn/. Briefly, the PDB and chain ID of PlzA was entered for the input receptor molecule to retrieve the crystal structure of the protein for the analysis. For each analysis, the corresponding nucleic acid sequence of interest was then pasted as the input ligand molecule, and the type of nucleic acid selected from the drop-down menu. The input nucleic acid sequences are listed in Table 2. All analyses were performed with default parameters. The top model for each computation is shown. Docking and confidence scores are provided in the corresponding figure legends. The more negative the docking score, the more probable the binding. A confidence score above 0.7 indicates the molecules would be highly likely to bind.

Protein structure prediction for PlzA_NTD_, PlzA_CTD_, and PlzA_RD-RD_ was performed via AlphaFold. All output PDB files generated from the various analyses were viewed with ChimeraX 1.4 (Goddard *et al*., 2018; Jumper *et al*., 2021; Pettersen *et al*., 2021).

### Cyclic di-nucleotide LC-MS analysis

Detection of cyclic di-nucleotides in recombinant protein purifications was conducted by liquid chromatography-tandem mass spectrometry (LC-MS/MS) at the Michigan State University Mass Spectrometry core. Briefly, recombinant proteins were purified and quantified as mentioned previously and subsequently dialyzed into EMSA buffer without Tween-20. For LC-MS/MS analysis, approximately 1 mg/mL of PlzA_WT_, PlzA_RD-RD,_ PlzA_NTD_, and PlzA_CTD_ were aliquoted and stored at −80°C until shipment to the core facility. Nucleotides bound to purified protein samples (1mg/mL) were analyzed by precipitating protein with 3 volumes of acetonitrile followed by centrifugation to pellet precipitated protein. The supernatant was transferred to a new tube and evaporated to dryness. The sample was reconstituted in mobile phase solvent A (10 mM tributylamine + 15 mM acetic acid in water/methanol, 97:3 v/v). LC-MS analysis was performed on a Waters Xevo G2-XS Quadrupole-Time-of-Flight (QTof) mass spectrometer interfaced with a Waters Acquity UPLC system. 10 µL of sample was injected onto a Waters Acquity BEH-C18 UPLC column (2.1×50 mm) and compounds were separated using the following gradient: initial conditions were 100% mobile phase A, hold at 100% A until 1 min, ramp to 99% B at 7 min (mobile phase B: methanol), hold at 99% B until 8 min, return to 100% A at 8.01 min and hold until 10 min. The flow rate through the column was 0.3 ml/min and the column temperature was held at 40°C. Compounds were ionized by electrospray ionization operated in negative mode. Capillary voltage was 2.0 kV, cone voltage was 35, source temperature was 100C and desolvation temperature was 350C. The cone gas and desolvation gas flows were 50 L/hr and 600 L/hr respectively. A TOF MS scan method with targeted enhancement of m/z 689 was used with 0.5 second scan time. Lockmass correction was performed using leucine enkephalin as the reference compound.

Data were processed using Waters Masslynx software. Extracted ion chromatograms were performed to look for the presence of c-di-GMP (m/z 689.09), c-di-AMP (m/z 657.09) and c-GAMP (673.09). The intensity values were converted to relative abundance as determined from the base peak values. The c-di-GMP concentration was determined from the peak area of the target compound against a standard curve of known c-di-GMP concentrations. These concentrations were calculated using the Targetlynx tool in the Waters Masslynx software. The percentage of protein bound with c-di-GMP was calculated from the total protein and calculated c-di-GMP concentrations in each sample.

### Circular dichroism spectroscopy

The CD analysis was performed at the University of Kentucky Center for Molecular Medicine Protein Core. Briefly, proteins were either buffer exchanged or diluted into 10 mM sodium phosphate buffer, pH 7.0. A 400 µL solution of each protein at a concentration of 100 µg/mL was pipetted into a 1 mm quartz cuvette and loaded into a Jasco J-810 spectrophotometer. The samples were scanned at 0.1 nm increments and CD spectra collected for wavelengths 185-260 nM with the temperature held constant at 25°C. A buffer only sample was used as a control. Two independent recombinant protein preparations were analyzed for each protein type. Spectral data was deconvoluted using CDtoolX. Deconvolution consisted of buffer spectra subtraction and the conversion of the CD spectra to molar residue ellipticity to normalize the data for sample concentration and sequence length. Data were visualized in GraphPad Prism 10.1.2.

Secondary structure prediction was performed using the BeStSel webserver using the single spectrum analysis function (Micsonai *et al*., 2015; Micsonai *et al*., 2018; Micsonai *et al*., 2022). Mean residue ellipticity values from the deconvoluted CD data were input into the server for analysis. The PDB file, 7MIE, was used to determine the secondary structure of the crystallized PlzA protein for comparisons to the generated recombinant proteins.

## AVAILABILITY

HDOCK (http://hdock.phys.hust.edu.cn/)

CDtoolX (https://cdtools.cryst.bbk.ac.uk/)

BeStSel (https://bestsel.elte.hu/index.php)

## ACKNOWLEDGEMENT

We dedicate this manuscript to our friend and colleague Christina R. Savage, Ph.D., whose initial studies on PlzA were invaluable to this work. We also thank Jessamyn Moore and Tatiana Castro Padovani for their support of the studies undertaken. Circular Dichroism was performed at the University of Kentucky Center for Molecular Medicine, and we thank Martin Chow for his assistance on these studies. The LC-MS/MS to detect di-nucleotides was performed by the Mass Spectrometry and Metabolomics Core at Michigan State University. A sincere thank you to Tony Schilmiller at MSU for all his help with the mass spectrometry services. Figures 1, 4a, 6a, 7a, 10a, and 13 were created via BioRender. Lastly, we thank Dr. József Kardos at Eötvös Loránd University for providing the secondary structure analysis for the 7MIE PDB file via email.

## Supplementary Information

**FIGURE S1.**
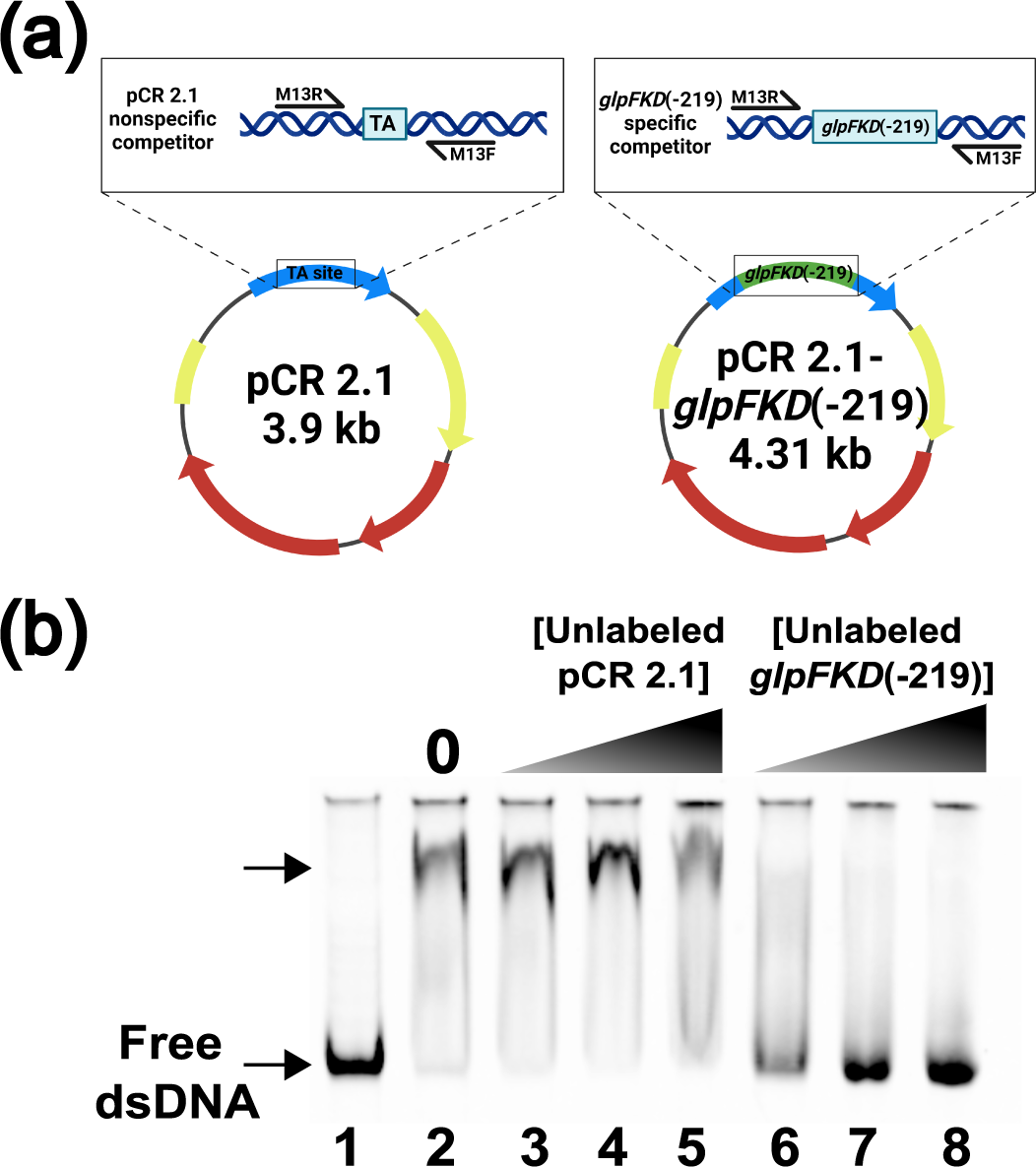
PlzA recognizes *glpFKD*(−219) and not pCR 2.1 sequences. (**a**) A schematic of the unlabeled DNA competitors derived from the empty pCR 2.1 vector backbone or the vector containing the *glpFKD*(−219) sequence. A labeled M13R and an unlabeled M13F primer were used to amplify the target DNAs from the respective plasmids to generate competitors. These competitors were then used as a control in competition EMSAs performed with labeled *glpFKD*(−219) DNA probe. (**b**) Lane 1 and lane 2 are probe only and probe + PlzA_WT_ controls respectively. All lanes contained a c-di-GMP concentration of 100 µM, a final poly-dI-dC concentration of 2.5 ng/µL, and a labeled *glpFKD*(−219) probe concentration of 50 nM. Lanes 2-8 contained a constant PlzA_WT_ protein concentration of 1.5 µM. The unlabeled pCR 2.1 competitor was titrated into the reactions at increasing molar excess relative to labeled probe: Lane 3-300 nM (6x), lane 4-500 nM (10x), and lane 5-1000 nM (20x). Unlabeled *glpFKD(*-219) competitor was titrated into the reactions at increasing molar excess relative to labeled probe: Lane 6-300 nM (6x), lane 7-500 nM (10x), and lane 8-600 nM (12x). The free arrow indicates PlzA-*glpFKD*(−219) complexes. The pCR 2.1 competitor could not compete away the PlzA-*glpFKD*(−219) complex at any tested concentration. Unlabeled *glpFKD*(−219) competitor was able to compete at all tested concentrations as low as 6x molar excess.

